# Comparative genomics and transcriptomics reveal differences in effector complement and expression between races of *Fusarium oxysporum* f.sp. *lactucae*

**DOI:** 10.1101/2024.04.11.589035

**Authors:** Helen J. Bates, Jamie Pike, R. Jordan Price, Sascha Jenkins, John Connell, Andrew Legg, Andrew Armitage, Richard J. Harrison, John P. Clarkson

## Abstract

This study presents the first genome and transcriptome analyses for *Fusarium oxysporum* f.sp. *lactucae* (Fola) which causes Fusarium wilt disease of lettuce. Long-read genome sequencing of three race 1 (Fola1) and three race 4 (Fola4) isolates revealed key differences in putative effector complement between races and with other *F. oxysporum* f.spp. following *mimp*-based bioinformatic analyses. Notably, homologues of *Secreted in Xylem* (*SIX*) genes, also present in many other *F. oxysporum* f.spp, were identified in Fola, with both *SIX9* and *SIX14* (multiple copies with sequence variants) present in both Fola1 and Fola4. All Fola4 isolates also contained an additional single copy of *SIX8*. RNAseq of lettuce following infection with Fola1 and Fola4 isolates identified highly expressed effectors, some of which were homologues of those reported in other *F. oxysporum* f.spp. including several in *F. oxysporum* f.sp. *apii*. Although *SIX8*, *SIX9* and *SIX14* were all highly expressed in Fola4, of the two *SIX* genes present in Fola1, only *SIX9* was expressed as further analysis revealed that copies of *SIX14* gene copies were disrupted by insertion of a transposable element. Two variants of Fola4 were also identified based on different genome and effector-based analyses. This included two different *SIX8* sequence variants which were divergently transcribed from a shared promoter with either *PSE1* or *PSL1* respectively. In addition there was evidence of two independent instances of HCT in the different Fola4 variants. The involvement of helitrons in Fola genome rearrangement and gene expression is discussed.

## 1 Introduction

*Fusarium oxysporum* is a globally important fungal species complex that includes plant pathogens, human pathogens, and non-pathogens (Edel-Hermann and Lecomte, 2019). Plant pathogenic isolates are grouped into different *formae speciales* (f.spp.) depending on their host range (generally one species) and are of major significance as they cause vascular wilts, crown and root rots of many important horticultural crops and ornamental plants (Edel-Hermann and Lecomte, 2019).

Lettuce (*Lactuca sativa*) is a globally significant vegetable crop, cultivated in over 150 countries worldwide (FAOSTAT) with a substantial market value of more than €3.3 billion in Europe (Eurostat, 2019) and $2 billion in the USA (United States Department of Agriculture - National Agricultural Statistics Service, 2021). However, lettuce production is becoming increasingly affected by Fusarium wilt disease caused by *F. oxysporum* f.sp. *lactucae* (Fola) which causes plant yellowing, stunting, wilting and death with losses of more than 50% commonly reported (Gilardi et al., 2017a). Fusarium wilt of lettuce was first described in Japan in 1955 (Motohashi, 1960) and since then has been identified in many lettuce producing areas of the world with an increasing number of outbreaks recently reported for the first time. Four races are described for Fola with race 1 (Fola1) the most established and widespread, primarily causing disease in field-grown lettuce in warmer parts of the world such as Asia, USA, Southern Europe and South America while races 2 and 3 are confined to Japan and Taiwan (Gilardi et al., 2017a). In contrast, Fola race 4 (Fola4) only emerged relatively recently in Northern Europe where Fusarium wilt of lettuce was previously completely absent and was first reported in 2015 affecting greenhouse-grown (protected) lettuce in the Netherlands (Gilardi et al., 2017a) and Belgium (Claerbout et al., 2018). Since then, Fola4 has been reported in other areas of Europe including the UK (Taylor et al., 2019b), Italy (Gilardi et al., 2019) and Spain (Gálvez et al., 2023). Notably, Fola4 initially only affected protected lettuce although has now also been identified in the open field in both Italy and Spain (Gilardi et al., 2019; Gálvez et al., 2023). In addition, Fola1 also seems to be spreading into Northern Europe with new reports on protected lettuce in both Norway (Herrero et al., 2021) and Northern Ireland (van Amsterdam et al., 2023). This suggests that in the near-future both Fola1 and Fola4 may co-exist in many locations affecting both protected and field-grown lettuce, as is currently the case in some European countries such as Italy (Gilardi et al., 2017b), Belgium (Claerbout et al., 2023) and Spain (Guerrero et al., 2020; Gálvez et al., 2023). Fola4 and Fola1 are very closely related based on phylogenetic analyses of DNA sequences of standard loci such as the translation elongation factor 1α, while Fola2 and Fola3 are in separate distinct clades (Gilardi et al., 2017a; Claerbout et al., 2023). Interestingly Fola4 has been shown to be more aggressive at lower temperatures than Fola1 (Gilardi et al., 2021). The four races of Fola can be distinguished based on their virulence on a differential set of lettuce cultivars (Gilardi et al., 2017a), which has recently been updated by the International Seed Federation (Worldseed.org). Moreover, there are also specific PCR assays published for both Fola1 (Pasquali et al., 2007) and Fola4 (Gilardi et al., 2017b). Although resistance to Fola1 is available within commercial cultivars (Garibaldi et al., 2004; Scott et al., 2010; Murray et al., 2021), these are not always adapted or suitable for some locations and are often also susceptible to Fola4. Moreover, the lettuce cultivars commonly used in indoor production in Europe are highly susceptible to Fola4 and hence the plant breeding industry has had to try and react quickly to identify and deploy resistant cultivars to this new race.

It is now well established that *F. oxysporum* has a compartmentalised genome consisting of both core and accessory (or lineage specific, LS) chromosomes with the former highly conserved and syntenous between different *F. oxysporum* f.spp. In contrast, the accessory chromosomes are highly variable, are dispensable for viability and are enriched with an abundance of transposable elements (Ma et al., 2010; Yang et al., 2020) which makes assembly of these regions difficult. It is also apparent through *F. oxysporum* genome analyses that some accessory chromosomes (referred to as pathogenicity chromosomes) contain the majority of effectors known to have a role in disease and are therefore associated with pathogenicity and host specificity (Vlaardingerbroek et al., 2016; van Dam et al., 2017; Armitage et al., 2018; Li et al., 2020; Ayukawa et al., 2021). An important finding initially for *F. oxysporum* f.sp. *lycopersici* (Fol) was that effectors were frequently found downstream of miniature impala elements (*mimps*), a specific family of miniature inverted-repeat transposable elements (MITEs) (Schmidt et al., 2013) and since then this has been used as a successful bioinformatics approach to identify the effector complement within other *F. oxysporum* f.spp. (van Dam et al., 2016; van Dam et al., 2017; van Dam and Rep, 2017; Armitage et al., 2018; Taylor et al., 2019a; Chang et al., 2020; Li et al., 2020). Interestingly, non-pathogenic *F. oxysporum* strains including the well-studied endophyte and biological control agent Fo47 (Aimé et al., 2013) also have accessory chromosomes and some effector genes, but are reported to have far fewer than pathogenic f.spp. (Constantin et al., 2021). Of the effectors identified in *F. oxysporum*, the most studied are the *Secreted in Xylem* (*SIX*) genes first identified in Fol which encode small, secreted proteins that are released into the xylem upon infection (Houterman et al., 2007). Since then, homologues of 14 Fol *SIX* genes have been identified in different numbers and complements in most *F. oxysporum* f.spp. and have therefore been used to differentiate between them based on presence / absence or sequence variation (Jangir et al., 2021). A further *SIX* gene, designated *SIX15* was identified in the Fol genome following comparison with the *F. oxysporum* f. sp. *physali* (infects cape gooseberry) genome (Simbaqueba et al., 2021). Moreover, *SIX* gene complement and sequence can also vary between races within a single *F. oxysporum* f.sp. For instance, the breaking of I gene-mediated resistance in tomato by Fol race 2 was shown to be associated with loss of the avirulence gene *SIX4* (*Avr 1*) or in the case of Fol race 3 by mutations in *SIX3* (*Avr 2*) (Takken and Rep, 2010). Similarly, in *F. oxysporum* f.sp. *cubense* (Focub) races affecting banana, *SIX1*, *SIX6*, *SIX9*, and *SIX13* were detected in race 1, *SIX1*, *SIX2*, *SIX7*, *SIX8*, and *SIX9* in race 4, while tropical race (TR) 4 carries *SIX1*, *SIX2*, *SIX6*, *SIX8*, *SIX9*, and *SIX13* (Czislowski et al., 2018). Three copies and four sequence variations of *SIX1* were also identified in Focub race 4 compared with one copy and two variants in race 1 (Guo et al., 2014). The *SIX8* sequence is also different between Focub races 4, TR4 and subtropical (STR4) races (Fraser-Smith et al., 2014). There is evidence that both horizontal chromosome transfer (HCT) and horizontal gene transfer (HGT) play a role in shuffling pathogenicity genes and chromosomes between different members of the *F. oxysporum* species complex (FOSC) potentially enabling host jumps and the emergence of new races within a f.sp. (Ma et al., 2010; Van Dam and Rep, 2017; Henry et al., 2020), although the mechanism for this is not clear.

Given the importance of Fola and the recent emergence of a new race that has expanded the pathogen’s geographic range, understanding the genetic differences between Fola1 and Fola4 and hence the implications for durability of resistance and evolution of the pathogen is of paramount importance.

## 2. Materials and methods

### Fusarium oxysporum isolates

The Fola and other *F. oxysporum* isolates used in this study were obtained from other researchers and industry collaborators or were directly isolated from host plants as described by Taylor et al., (2019c) (Table 1). Briefly, this involved surface sterilising sections of symptomatic stem or tap root tissue that exhibited typical vascular browning in 70% ethanol followed by washing twice in sterile distilled water (SDW) and placing on potato dextrose agar (PDA) containing 20 µg/ml of chlorotetracycline. Plates were incubated for 4 days at 20°C, from which emerging *Fusarium* colonies were then sub-cultured from hyphal tips onto fresh PDA. Spore suspensions of all isolates were prepared from two-week-old PDA cultures grown at 25°C in potato dextrose broth (PDB) amended with 20% glycerol (v/v) for long-term storage on ceramic beads at −80°C. In total, three Fola1 isolates (AJ520, AJ718, AJ865) and three Fola4 isolates (AJ516, AJ592, AJ705) were selected as the focus of this study and were subject to genome sequencing, effector identification and RNAseq *in planta* alongside an isolate of *F. oxysporum* f.sp *matthiolae* (Foma isolate AJ260) which infects column stocks (*Matthiola incana*). Isolates of *F. oxysporum* infecting *Narcissus* spp. (*F. oxysporum* f.sp. *narcissi*; Fon isolate AJ275 [FON63], Taylor *et al*., 2019a) and wild rocket (*Diplotaxis tenuifolia*; *F. oxysporum* rocket isolate AJ174, Taylor *et al*., 2019b) were used for genome sequencing and comparison of effector complement only. The non-pathogenic *F. oxysporum* endophyte isolate Fo47 used in other genome studies (e.g. Armitage *et al*., 2018, van Dam *et al*., 2016) was also used for comparative purposes and RNAseq. All isolates were confirmed as *F. oxysporum* by PCR and sequencing of the translation elongation factor (TEF) gene as described by Taylor *et al*., (2016). Fola isolate race identity and virulence was confirmed by testing on a standard lettuce differential set (Gilardi *et al*., 2017a) by isolate suppliers or in previous studies and also by confirming the presence of *SIX9* and *SIX14* in Fola1 and *SIX8*, *SIX9* and *SIX14* in Fola4 isolates by PCR as described by Taylor et al. (2016) using modified primers for Fola (Supplementary data SD1-T1). These different complements of *SIX* genes for Fola1 and Fola4 were consistent across multiple Fola isolates in initial studies (unpublished). The Foma and *F. oxysporum* rocket isolates were confirmed to be pathogenic on their respective hosts using standard root dip inoculations with conidial suspensions as described by Taylor et al., (2019b) while Fon isolate AJ275 was previously confirmed to be pathogenic on daffodil bulbs (Taylor et al., 2019a).

**Table 1.** *Fusarium oxysporum* f.sp. *lactucae* and other *F. oxysporum* isolates used in this study.

| Isolate code | Identity | Race | Host source | Country of origin | Other code / Reference / Source |
| --- | --- | --- | --- | --- | --- |
| AJ520 | *F. oxysporum* f.sp. *lactucae* | 1 | Lettuce (*Lactuca sativa*) | Italy | ATCCMya-3040<br>Gilardi et al., 2017 |
| AJ718 | *F. oxysporum* f.sp. *lactucae* | 1 | Lettuce (*Lactuca sativa*) | France | Fyto 7211<br>BASF, NL |
| AJ865 | *F. oxysporum* f.sp. *lactucae* | 1 | Lettuce (*Lactuca sativa*) | Spain | PF1<br>Gs Espana, SP |
| AJ516 | *F. oxysporum* f.sp. *lactucae* | 4 | Lettuce (*Lactuca sativa*) | England | LANCS1<br>Taylor et al., 2019c |
| AJ592 | *F. oxysporum* f.sp. *lactucae* | 4 | Lettuce (*Lactuca sativa*) | England |  |
| AJ705 | *F. oxysporum* f.sp. *lactucae* | 4 | Lettuce (*Lactuca sativa*) | Netherlands | AD035<br>Enza Zaden, NL |
| AJ260 | *F. oxysporum* f.sp. *matthiolae* | - | Column stocks (*Matthiola incana*) | England |  |
| AJ275 | *F. oxysporum* f.sp. *narcissi* | - | Daffodil (*Narcissus pseudonarcissus*) | England | FON63<br>Taylor et al., 2019a |
| AJ174 | *F. oxysporum* | - | Rocket (*Diplotaxis tenuifolia*) | England | Taylor et al., 2019b |
| AJ937 | *F. oxysporum* | - | Endophyte | France | Fo47, Aimé et al, 2013 |

### Genome sequencing, assembly and analyses

#### DNA extraction

*Fusarium oxysporum* isolates (Table 1) were grown on PDA plates at 25°C for 4 days and 5 mm mycelial plugs from growing tips then used to inoculate 50 ml potato dextrose broth (PDB) containing 20 µg/ml streptomycin which were incubated for 4 days in the dark at 25°C at 180 rpm. The resulting mycelium for each isolate was harvested on filter paper (Fisherbrand 11556873, Fisher Scientific UK Ltd.), washed with distilled water, blotted dry and snap frozen in liquid nitrogen prior to freeze drying and storing at −80°C. Freeze-dried mycelium (20 mg) was used for DNA extraction using a Macherey-Nagel NucleoSpin Plant II kit (Fisher Scientific) following the manufacturer’s instructions. DNA samples were eluted in 30 µl 10 mM Tris HCl pH 8 and analysed for purity and quantity using the Nanodrop spectrophotometer (NanoDrop One, ThermoFisher Scientific) and Qubit fluorometer (Invitrogen) and for integrity using the Agilent TapeStation (TS 4150, Agilent Technologies). DNA samples with Nanodrop ratios 260/280 between 1.8-2.0, 260/230 between 2.0-2.2 and molecular weight >50 kb were selected for sequencing.

#### Genome sequencing

Illumina PCR-free genome sequencing was carried out for isolates Foma AJ260, Fon AJ275, Fola1 AJ520 and Fola4 AJ516 as previously described (Armitage et al., 2018). Long-read genome sequencing was carried out using 1 µg input DNA into Oxford Nanopore Technologies (ONT) ligation sequencing kit LSK108 (Foma AJ260, Fon AJ275) or LSK110 (Fola1 isolates AJ520, AJ718, AJ865; Fola4 isolates AJ516, AJ592, AJ705 and *F. oxysporum* rocket) as described and flow cells FLO-MIN106 R9.4 (Foma AJ260, Fon AJ275) or FLO-MIN106 R9.4.1 (all other isolates). Enzymes were purchased from New England Biolabs (NEBNext Companion Module #E7180) and Omega Bio-Tek Mag-bind TotalPure NGS magnetic beads were from VWR International. All ONT sequencing runs were performed on the GridION with live high-accuracy base calling. Data were re-base called using super-high accuracy mode post-run and prior to genome assembly.

#### Genome assembly

*De novo* genome assemblies for the six Fola (3 x Fola1, 3 x Fola4), and the single isolates of Foma, and *F. oxysporum* rocket were generated from long-read Oxford Nanopore Technologies sequence data (Wang et al., 2021). Quality control of ONT data was performed using NanoPlot v1.30.1 (De Coster et al., 2018). Adapter trimming was performed using Porechop v0.2.4 (https://github.com/rrwick/Porechop) with default parameters, followed by removal of reads shorter than 1 kb or with a quality score less than Q9 using Filtlong v0.2.1 (https://github.com/rrwick/Filtlong). Long read data were assembled using NECAT v0.0.1_update20200803 (Chen et al., 2021) with a genome size of 60 Mb, with other parameters left as default. Long read error correction was performed by aligning reads to the assemblies with Minimap2 v2.17-r941 (Li, 2018) to inform one iteration of Racon v1.4.20 (Vaser et al., 2017), followed by one iteration of Medaka v1.5.0 (https://github.com/nanoporetech/medaka) using the r941_min_high_g360 model. Quality control of Illumina paired-end reads was performed using FastQC v0.11.9 (https://www.bioinformatics.babraham.ac.uk/projects/fastqc/), with adapters and low-quality regions trimmed using Fastq-Mcf v1.04 (Aronesty, 2013). Bowtie2 v2.2.5 (Langmead & Salzberg, 2012) and SAMtools v1.13 (Li et al., 2009) were used to align short reads to the long-read assemblies. Three iterations of polishing were carried out using Pilon v1.24 (Walker et al., 2014) to allow correction of single base call errors and small insertions or deletions. Assembly statistics were generated using a custom Python script, with assembly quality also assessed through single copy ortholog analysis performed using BUSCO v5.2.2 (Simão et al., 2015), with the hypocreales_odb10 database. Genome assembly of the Fon isolate AJ275 was performed prior to this study using the pipeline described in Armitage et al. (2020).

### Gene prediction in Fola1, Fola4 and Foma isolates

Prior to gene prediction repetitive sequences were masked using RepeatMasker version 4.1.2 (Smit et al., 2013-2015) and RepeatModeler version 2.0.5 (Smit and Hubley, 2008-2015) to produce both hard and soft masked genome assemblies. Gene models were annotated using both BRAKER version 2.16 (Brůna *et al*., 2021) and CodingQuarry version 2 gene prediction tools (Testa *et al*., 2015). RNAseq reads (see below) were adapter trimmed and poor-quality reads removed using Fastq-Mcf version 1.04 (Aronesty, 2013). Pre-processed reads were aligned to respective *F. oxysporum* assemblies using STAR version 2.7.10 (Dobin *et al*., 2013) with the flags --winAnchorMultimapNmax set to 200 and --seedSearchStartLmax set to 30 to improve mapping sensitivity. An initial round of gene prediction was performed using the BRAKER 2 method with flags --fungus to specify a fungal organism, and -- softmasking to specify a soft masked genome assembly. RNAseq alignments generated for Fola infection of lettuce (as described below) were used as inputs to BRAKER to provide additional evidence to improve gene prediction accuracy. A second round of gene prediction was performed using the CodingQuarry pathogen method. Genome alignment files produced by STAR as described above were used as inputs to Cufflinks version 2.2.1 (Trapnell *et al*., 2010) to produce a *de novo* transcriptome which was used as a guide for CodingQuarry. Gene predictions for both BRAKER and CodingQuarry were investigated using the BEDTools intersect function (Quinlan & Hall, 2010) with the -v flag to separate CodingQuarry predictions where no overlap to a BRAKER prediction was found. Resulting gene predictions were combined and ordered with any duplicated annotations removed using a custom-made Python script to retain BRAKER gene models and integrate CodingQuarry predictions that were located in intergenic regions (Python Software Foundation, 2021). Following this, predicted genes were functionally annotated using InterProScan version 5.62-94.0 (Jones *et al*., 2014). Orthologous gene families were identified using OrthoFinder version 2.5.4 (Emms & Kelly, 2019) with the Fola1 (AJ520, AJ718, AJ865), Fola4 (AJ516, AJ592, AJ705), Foma AJ260 genomes, along with the *F. oxysporum* Fo47 genome.

### Identification and visualisation of genomic features for Fola1 and Fola4 isolates

Annotation of genomic features including transposable elements, carbohydrate active enzymes (CAZymes), fungal effector proteins, secreted signal peptides and secondary metabolite clusters was carried out for Fola1 and Fola4 isolates. Transposable elements were identified using transposonPSI version 1.0.0 (Haas, 2010) with the default settings. CAZymes were identified using the hmmscan tool from HMMER version 3.3.2 (Finn *et al*., 2011) with HMM models from the dbCAN2 version 11 database (Zhang *et al*., 2018). Candidate fungal effector proteins were identified using EffectorP version 3.0 (Sperschneider *et al*., 2016) with default settings. Secreted signal peptides were predicted by taking a unique list of genes from the combined output of SignalP versions 2, 3, 4 and 5 (Almagro Armenteros *et al*., 2019, Bendtsen *et al*., 2004, Nielsen & Krogh, 1998, Petersen *et al*., 2011). Secondary metabolite clusters were identified using the antiSMASH version 6.0 webserver (Blin *et al*., 2021) with the addition of the cluster-border prediction based on transcription factor binding sites feature enabled. Individual secondary metabolite genes were pulled out from the genebank file produced by antiSMASH using a custom python script. Visualisation of core and pathogenicity chromosomes was performed for Fola1 AJ520 and Fola4 AJ516 using Circos software version 0.69-9 (Krzywinski *et al*., 2009). The locations of genomic features including CAZymes, transposable elements, secondary metabolite clusters, miniature impala sequences, and *SIX* gene homologs that were identified as described above, were plotted on the ideogram based on their genomic position. The putative effector gene locations generated by the *mimp*-associated pipeline described below, were mapped to the annotated genome to identify genes and those genes filtered by EffectorP and SignalP predictions to produce the list of putative effector genes shown. To visualise reads that map to both Fola1 AJ520 and Fola4 AJ516 genomes from the other *F. oxysporum* isolates, short read data was simulated from the long reads for each genome by generating 300 bp fragments with a 50 bp step. This simulated short read data for Fola1 isolates AJ718 and AJ865, Fola4 isolates AJ592 and AJ705, Foma AJ260 and the non-pathogenic *F. oxysporum* Fo47 were aligned using BWA-MEM version 0.7.17-r1188 (Li, 2013) using default parameters. Genome alignment coverage was calculated using BEDTools genomecov function (Quinlan & Hall, 2010) with the flag -d to specify that a per base coverage should be calculated. A custom python script was employed to calculate a per window sum of genome coverage at a window size of 10,000 bp.

### Phylogenetic reconstruction of *F. oxysporum* core and accessory genomes

To investigate the genetic relatedness of core and accessory genome sequences for the *F. oxysporum* genomes included in this study (Supplementary data SD1-T2), alignment-free phylogenetic reconstruction was performed using SANS ambages v2.3_9A (Rempel and Wittler 2021). Highly contiguous genome sequences were aligned to the core chromosomes of the reference *F. oxysporum* f.sp. *lycopersici* isolate 4287 using Minimap2 v2.17-r941 (Li, 2018). Contigs with continuous alignments of more than 100 kb were then extracted as core genome sequences. To prevent interference from repetitive and unplaced contigs, sequences less than 100 kb were removed from the remaining contigs following extraction of core genome sequences. The remaining contigs were considered as accessory genome sequences. Using these core and accessory sequences, SANS ambages v2.3_9A was run with 1000 bootstrap replicates and the option ‘strict’ to output a Newick format file, and all other options left as default. Phylogenetic trees were visualised using iTOL v6 (Letunic and Bork, 2021).

### Synteny of Fola core and accessory genomes

To compare the synteny across core and accessory genomes of all the Fola1 and Fola4 isolates, collinearity analysis was performed using the core and accessory contigs extracted for phylogenetic reconstruction as described above. Pairwise alignments were performed using Minimap2 v2.17-r941 (Li, 2018) and prior to visualisation, core genome alignments smaller than 100 kb and accessory genome alignments smaller than 10 kb were discarded. The results were visualised using NGenomeSyn v1.41 (He et al., 2023).

### *Mimp*-associated putative effector identification in Fola and other *F. oxysporum* genome assemblies

As highlighted previously, miniature impala elements (*mimps)* have been used previously to predict potential effectors in *F. oxysporum* including the majority of *SIX* genes in different *F. oxysporum* f.spp. (Armitage et al., 2018, Brenes Guallar *et al*., 2022, van Dam et al., 2016, van Dam & Rep, 2017, Schmidt *et al*., 2013). We therefore adapted a *mimp*-associated effector identification pipeline as utilised by van Dam *et al*., (2016), to find putative effectors in genomes of all the Fola isolates and other *F. oxysporum* f.spp. generated in this study (Fon AJ275, Foma AJ260, and *F. oxysporum* rocket AJ174). Alongside these assemblies, publicly available representative high-quality genomes (minimum ≤50 contigs and a reported BUSCO of >97%) for the *F. oxysporum* endophyte Fo47, and the *F. oxysporum* f.spp. *cepae, conglutinans, coriandrii, cubense, lini, lycopersici, niveum, rapae* and *vasinfectum* were downloaded from GenBank (https://www.ncbi.nlm.nih.gov/data-hub/genome/) (Supplementary data SD1-T2).

Two methods of searching for *mimps* were employed for each *F. oxysporum* genome. The first used a custom python script to search for the *mimp* TIR sequences, "CAGTGGG..GCAA[TA]AA" and "TT[TA]TTGC..CCCACTG". Where sequences matching this pattern occurred within 400 nucleotides of each other a *mimp* was recorded. The second *mimp* searching method employed a Hidden Markov Model (HMM) which was developed using the HMM tool HMMER (3.3.1) (Wheeler & Eddy, 2013). Briefly, publicly available *mimp* sequences (Supplementary data SD1-T3) and further sequences identified using the regular expression method in the *Fusarium* assemblies were used to build a *mimp* profile-HMM. This profile-HMM was used as the input for an NHMMER search of each genome. Using *mimps* identified by both *mimp* finding methods, sequences 2.5 kb upstream and downstream of each *mimp* were extracted and subjected to gene prediction using AUGUSTUS (3.3.3) (Stanke *et al*., 2006) with the “*Fusarium*” option enabled, and open reading frames (ORFs) within this region identified using the EMBOSS (6.6.0.0) tool, getorf (https://www.bioinformatics.nl/cgi-bin/emboss/getorf). ORFs and gene models from each genome with a signal peptide predicted using SignalP (4.1, default settings) (Petersen et al., 2011) were then clustered using CD-HIT (4.8.1) (Fu *et al*., 2012) to create a non-redundant candidate effector set for each genome. Sequences of between 30 aa and 300 aa were then extracted from the individual, non-redundant candidate effector sets for all isolates and were combined and clustered using CD-HIT (4.8.1) at 65% identity. The longest sequence from each cluster was then subject to effector prediction using EffectorP (2.0.1) (Sperschneider et al., 2016), generating a collective candidate *mimp*-assoicted-effector set. To determine the presence/absence of the candidate effectors across all of the *Fusarium* genomes, the collective candidate *mimp*-associated-pan-effector set was used to search for effector homologues across all genomes using TBLASTN, with a cut-off 1e^-6^ and a percentage identity and coverage threshold of 65%. TBLASTN hits were extracted, translated using transeq from EMBOSS (6.6.0.0), and filtered by the presence of a signal peptide (SignalP version 1.4, default settings) and effector prediction using EffectorP (2.0.1, default settings). Candidates were classified into effector groups at 65% identity CD-HIT (4.8.1) and a presence/absence data matrix was generated using the candidate effector clusters. The R functions dist(method = binary) and hclust were used to cluster the binary effector matrix and the R package Pheatmap (1.0.12) (Kolde & Kolde, 2015) used to create the heatmap (R version 3.6.3).

### Identification of *SIX* genes in Fola and other *F. oxysporum* genome assemblies

The complement of *SIX* gene homologues was determined in all the Fola and other *F. oxysporum* isolate genomes generated in this study as well as in publicly available genomes for the *F. oxysporum* endophyte Fo47, and for *F. oxysporum* f.spp. *capsici, cepae, conglutinans, coriandrii, cubense, lini, luffae, lycopersici, matthiolae, niveum, rapae* and *vasinfectum* which were downloaded from GenBank following a genome search (https://www.ncbi.nlm.nih.gov/data-hub/genome/) (Supplementary data SD1-T2). Reference sequences for *SIX1-SIX15* from *F. oxysporum* f.sp *lycopersici* (isolate 4287) were downloaded from the NCBI database (Supplementary data SD1-T4) and homologues of each *SIX* gene identified in each assembly using tBLASTx (1e-6 cut-off). A binary data matrix indicating presence (“+”) or absence (“-”) was generated using the tBLASTx hit data. *SIX* gene phylogenies were then constructed for the *SIX* genes present in Fola1 (*SIX9*, *SIX14*) and Fola4 (*SIX8*, *SIX9*, *SIX14*). The locations of *SIX8*, *SIX9*, and *SIX14* tBLASTx (1e-6 cut-off) hits were recorded, and the sequence within this region extracted using Samtools (version 1.15.1). Extracted regions from each genome were added to a multiFASTA file for each *SIX* gene. MAFFT (version 7.505) (Katoh *et al*., 2019) was used to construct a multiple sequence alignment using the “—adjustdirectionaccurately” and “–reorder” options. To ensure correct alignment, any overhanging regions were inspected and trimmed manually. IQ-TREE (Version 2.2.0.3) (Nguyen *et al*., 2015) was used to infer a maximum-likelihood phylogeny using the ultrafast bootstrap setting for 1000 bootstrap replicates and was visualised using iTOL (Letunic & Bork, 2021).

### Expression of predicted Fola effectors in lettuce

#### Inoculation of lettuce seedlings with Fola and non-pathogenic *F. oxysporum*

Lettuce seedlings were grown and inoculated with isolates of Fola1 AJ520 and two variants of Fola4 (AJ516, AJ705) on an agar medium using a method adapted from Taylor et al. (2016) for subsequent RNA extraction and sequencing. As controls, two *F. oxysporum* isolates non-pathogenic on lettuce were also used for inoculations; these were Foma AJ260 which is pathogenic on column stocks where the genome contains homologues of *SIX8* and *SIX9* also both present in Fola4, and the endophyte *F. oxysporum* Fo47 which is a ‘standard’ non-pathogenic isolate used in other studies (Constantin *et al*., 2021) where the genome contains no *SIX* genes. A non-inoculated control treatment (no *F. oxysporum*) was also set up. Autoclaved ATS medium (5 mM KNO_3_, 2.5 mM KPO_4_, 3 mM MgSO_4_, 3 mM Ca(NO_3_)^2^, 50 uM Fe-EDTA, 70 uM H_3_BO_3_, 14 uM MnCl_2_, 0.5 uM CuSO_4_, 1 uM ZnSO_4_, 0.2 uM Na_2_MoO_4_, 10 uM NaCl, 0.01 uM CoCl_2_, 0.45% Gelrite (Duchefa Biochemie, Haarlem, The Netherlands)) was used to three-quarter fill square petri dishes (12 x 12 x 1.7 cm, Greiner Bio-One, UK) and once set, the top 5 cm of the gel was removed with a sterile spatula.

Lettuce seeds of a Fola susceptible cultivar (cv. Kordaat) were surface sterilised in a 10% bleach water (v/v) solution for 5 min and then rinsed three times with SDW before they were placed across the cut edge of the agar in each plate (12 seeds per plate) after which the lid was replaced and secured with tape. Stacks of five plates were wrapped in cling film and placed at 4°C in the dark for 4 days. Plates were then incubated at 15°C in 16 h light/dark for 8 days, before the temperature was increased to 25°C for six days. Conidial suspensions of each *F. oxysporum* isolate were prepared by releasing spores from two-week-old cultures grown on PDA at 25°C with 10 mL SDW and filtering through three layers of Miracloth. Spore suspensions were adjusted to 1 x 10^6^ spores ml^-1^ in SDW with the addition of 200 µl of Tween20 l^-1^ and 1.5 ml pipetted directly onto the lettuce roots on each plate and spread by tilting, before being dried briefly under sterile air flow. Non-inoculated control treatments consisted of SDW + Tween only. In total, seven agar plates of lettuce seedlings were inoculated for each *F. oxysporum* isolate, of which four were used for RNAseq and three were incubated further to confirm that disease developed for the Fola isolates and not for the non-pathogenic Foma AJ260 and the *F. oxysporum* endophyte isolate Fo47. Plates were placed in a randomised design within an incubator at 25°C (16 h photoperiod). Lettuce roots from each treatment for RNAseq were harvested 96 hours post-inoculation and pooled into one sample of 12 plants per plate. This time point corresponded to a peak in expression of *SIX8*, *SIX9* and *SIX14* as quantified through RT-PCR assays of a time-course study using the same bioassay for lettuce seedlings cv. Temira inoculated with Fola4 AJ516 (data not shown). Roots were rinsed in SDW, blotted dry and flash frozen in liquid nitrogen and stored at −80°C prior to RNA extraction and sequencing. Lettuce seedlings from each treatment retained at 25°C for disease assessment were inspected for root browning and death two weeks post-inoculation. To provide data for gene expression *in vitro* in order to identify upregulated genes *in planta*, spore suspensions of each *F. oxysporum* isolate (Fola1 AJ520, Fola4 AJ516 / AJ705, Foma AJ260 and *F. oxysporum* Fo47) were prepared as above, and 500 µL of 1 x 10^6^ spores ml^-1^ pipetted onto a PDA plate containing an autoclaved cellulose disc placed on the surface and incubated for 96h at 25°C. Six replicate plates were prepared and the resulting mycelium harvested by scraping off the layer growing on the cellulose surface before immediately flash freezing. Four replicate samples were then used for RNA extraction.

### RNA extraction and sequencing

Lettuce roots were ground to a fine powder using a pestle and mortar filled with liquid nitrogen and approximately 100 mg of tissue transferred to a 2 ml tube. Frozen root material was ground further using a Dremel drill (model 398, with a rounded drill bit) and then RNA extracted using Trizol ® reagent (Thermo Fisher Scientific) following the manufacturer’s guidelines. Extracted RNA was precipitated using 900 µl of lithium chloride to 100 µl RNA (250 µl LiCl2 + 650 µl DEPC treated water) and any DNA was removed from samples using DNase 1 (Sigma-Aldrich, UK). RNA samples were visualised on a 2% agarose gel (containing GelRedTM at 2 µl per 100 ml of gel) with the addition of loading dye (Orange G, Sigma-Aldrich, UK) to check for degradation. RNA samples (four replicates for each isolate) were sent to Novogene for polyA-enrichment, followed by Illumina PE150 sequencing at a depth of 23 Gb raw data per sample for *in planta* samples and 9 Gb per sample for *in vitro* mycelial RNA samples.

### RNAseq data analyses

RNAseq reads containing transcripts from both *F. oxysporum* isolates and lettuce were trimmed using Fastq-Mcf as described above. To separate *F. oxysporum* reads from lettuce, the lettuce genome (*Lactuca sativa*, NCBI accession: GCF_002870075.3) was downloaded from NCBI and used for alignment of pre-processed RNAseq reads using STAR as described above with the addition of the flag --outReadsUnmapped which was used to specify that a file of non-mapping reads should be created. Non-mapping putative *F. oxysporum* RNAseq reads were pseudo-aligned to the respective reference genome and quantified using Kallisto version 0.48.0 (Bray *et al*., 2016). Differential gene expression analysis was conducted using R version 4.1.3 (R core team, 2021) with the DESeq2 package version 1.34.0 (Love *et al*., 2014) using the contrast function to give a list of differentially expressed genes.

Results were filtered to identify putative effectors that were upregulated *in planta*. Firstly, all genes upregulated *in planta* with a log 2 fold change (L2FC) greater than 2.0 (compared to PDB grown controls) were identified for each isolate. These were then sorted for genes containing a signal peptide and with gene length less than 1 kb to identify small, secreted proteins. Genes identified as CAZymes, secondary metabolites, transposons or those with homologues of greater than 70% identity in the non-pathogen Fo47 were discarded from the analysis. Remaining genes were then sorted based on level of induction based on L2FC, identification as a putative effector from the previously described effector discovery pipeline and presence on an accessory contig. Expressed candidate effectors were grouped by orthogroup (supplementary data SD1-T12) and all members of the orthogroup were cross-referenced for level of expression. Putative effectors by orthogroup were also cross-referenced to gene locations of candidate effector clusters (CECs) identified by the effector discovery pipeline.

## 3. Results

### Core and accessory genome phylogenies support a single Fola1 and Fola4 clade

Nanopore sequencing produced high quality and highly contiguous genome assemblies for Fola, Foma, Fon and *F. oxysporum* rocket isolates (supplementary data SD1-T6). To identify core and accessory genome sequences, these *de novo* genome assemblies and selected high quality genome sequences from other *F. oxysporum* f. spp. (supplementary data SD1-T2) were aligned to the reference *F. oxysporum* f.sp. *lycopersici* isolate 4287 genome. Core genome contigs comprised approximately 45-55 Mb across all isolates (Table 2), consistent with observations in other *F. oxysporum* f.spp. Following removal of contigs less than 100 kb to prevent confounding results from unplaced core sequences and/or highly repetitive contigs, the size of the accessory genomes varied considerably across all the *F. oxysporum* isolates, from 1.3 Mb for *F. oxysporum* f.sp. *cubense* 16052 to 23.5 Mb for *F. oxysporum* f.sp. *lini* 39 (Table 2). The putative accessory genomes of all Fola1 isolates had similar sizes of around 15 Mb, while the Fola4 isolates had larger accessory genome sizes, with 21 Mb for AJ516 and approximately 18 Mb for AJ705 and AJ592.

**Table 2.** Number and sizes of core and accessory *Fusarium oxysporum* (*Fo*) genome sequences used for phylogenetic reconstruction.

| *F. oxysporum* f.sp. / isolate | Core genome |  | Accessory genome |  |
| --- | --- | --- | --- | --- |
|  | Contigs | Size (Mb) | Contigs | Size (Mb) |
| *Fo* f.sp. *apii* 207.A | 12 | 46.1 | 26 | 18.3 |
| *Fo* f.sp. *cepa*e Fus2 | 21 | 45.5 | 8 | 7.2 |
| *Fo* f.sp. *conglutinans* Fo5176 | 14 | 54.8 | 4 | 12.0 |
| *Fo* f.sp. *coriandrii* 3-2 | 9 | 46.3 | 22 | 15.5 |
| *Fo* f.sp. *cubense* 16052 | 11 | 49.9 | 1 | 1.3 |
| *Fo* f.sp. *cubense* UK0001 | 11 | 47.1 | 2 | 1.4 |
| *Fo* f.sp. *lactuca*e race4 AJ516 | 14 | 47.5 | 17 | 20.9 |
| *Fo* f.sp. *lactuca*e race1 AJ520 | 17 | 45.2 | 14 | 15.5 |
| *Fo* f.sp. *lactuca*e race4 AJ592 | 19 | 46.2 | 11 | 18.2 |
| *Fo* f.sp. *lactuca*e race4 AJ705 | 17 | 46.5 | 12 | 17.7 |
| *Fo* f.sp. *lactuca*e race1 AJ718 | 20 | 46.3 | 12 | 15.2 |
| *Fo* f.sp. *lactuca*e race1 AJ865 | 20 | 45.6 | 11 | 15.7 |
| *Fo* f.sp. *lini* 39 | 11 | 45.8 | 8 | 23.5 |
| *Fo* f.sp. *lycopersici* 4287 | 12 | 45.8 | 13 | 14.5 |
| *Fo* f.sp. *matthiolae* AJ260 | 11 | 46.0 | 23 | 13.9 |
| *Fo* f.sp. *narcissus* FON63 | 12 | 44.6 | 19 | 14.8 |
| *Fo* f.sp. *niveum* 110407-3-1-1 | 24 | 46.5 | 6 | 3.1 |
| *Fo* f.sp. *vasinfectum* TF1 | 12 | 45.9 | 3 | 4.0 |
| *Fo* rocket AJ174 | 19 | 50.4 | 10 | 12.1 |
| *F. oxysporum* Fo47 | 11 | 46.1 | 1 | 4.3 |

Phylogenies based on the identified core and accessory genome sequences were generated using an alignment free reconstruction method (Figure 1). Both phylogenies demonstrated a single well supported clade for all the Fola isolates, suggesting a monophyletic origin of Fola1 and Fola4. The topology of the Fola clade in the core and accessory trees were congruent, with Fola1 and Fola4 as sister clades. Within each Fola race there was further separation from a common ancestor into two groups. Within the Fola1 clade, AJ718 represented a separate group to AJ520 and AJ865, while in the Fola4 clade AJ516 was separated from AJ705 and AJ592 (Figure 1). Interestingly, Foma AJ260 was observed to share the closest common ancestor with the Fola clade in both the core genome (Figure 1A) and the accessory genome (Figure 1B) phylogenies, suggesting that the Fola clade shares a common ancestor with Foma AJ260.

**Figure 1.**
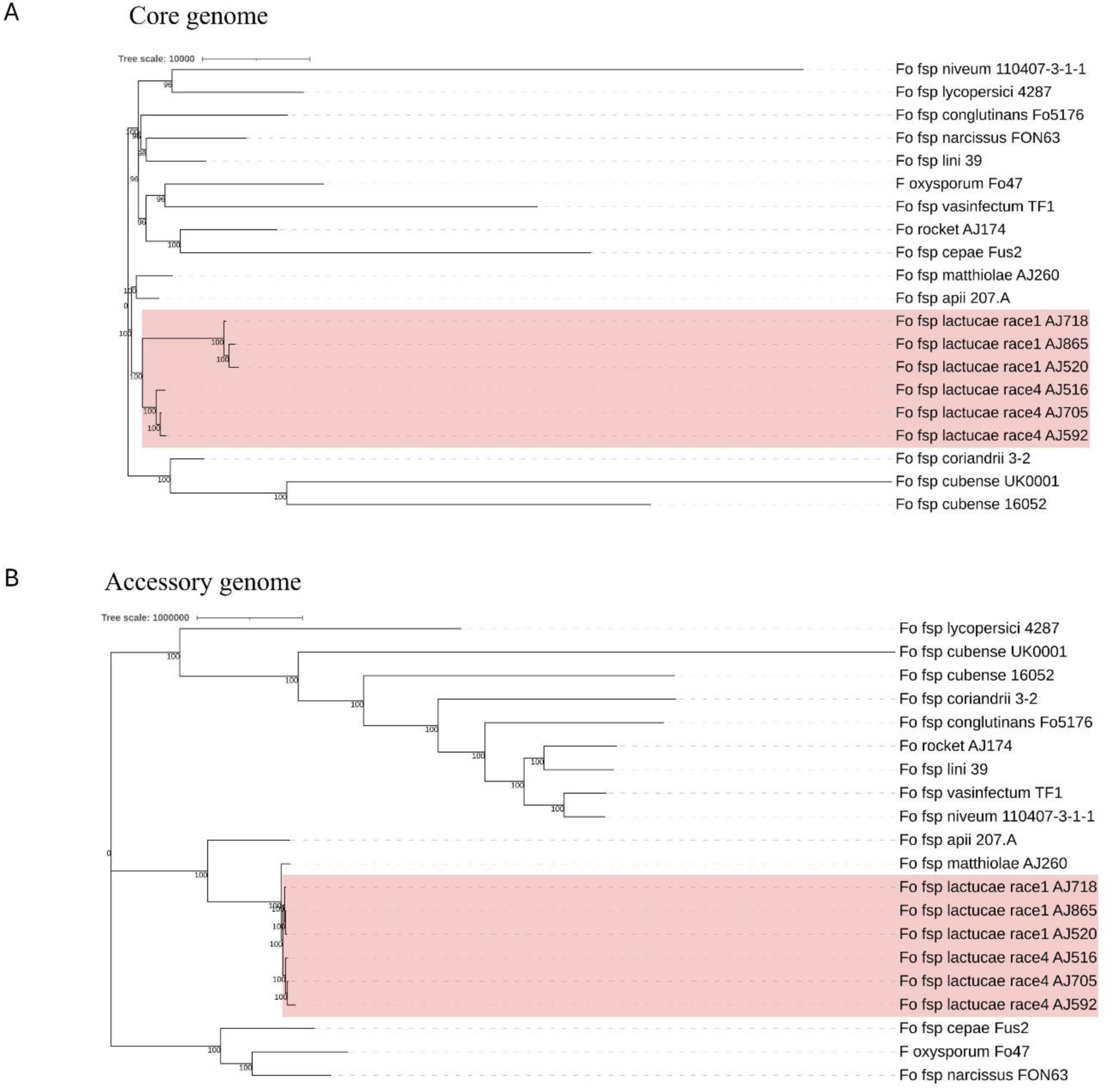
Core (A) and accessory (B) genome phylogenies for *Fusarium oxysporum* f. sp. *lactucae* (Fola) and other *F. oxysporum* f.spp. The monophyletic Fola clades are highlighted in the red boxes. Numbers at nodes represent percentage bootstrap support following 1000 replicates. The scale represents SNP count.

### Fola1 and Fola4 isolates cluster separately based on *mimp*-associated candidate effector profiles

To investigate differences in effector repertoire between Fola1 and Fola4, we used a *mimp*-based effector discovery pipeline to analyse six Fola genomes (3 x race 1 and 3 x race 4), along with 15 other *F. oxysporum* f.spp. with high quality genome assemblies (Table 3). The total number of candidate effectors across all these *F. oxysporum* genomes ranged from 52 in *F. oxysporum* f.sp. *cubense* race 1 isolate 160527 to 277 in Fola4 isolate AJ592 (Table 3). Interestingly the *F. oxysporum* (non-pathogenic) endophyte isolate Fo47 yielded 66 candidate effectors, which was 14 more than for *F. oxysporum* f. sp. *cubense* isolate 160527. Fewer candidate effectors were identified in the three Fola1 isolates compared with the three Fola4 isolates with 215, 230, 247 and 274, 276, 277 candidate effectors found, respectively (Table 3).

**Table 3.**
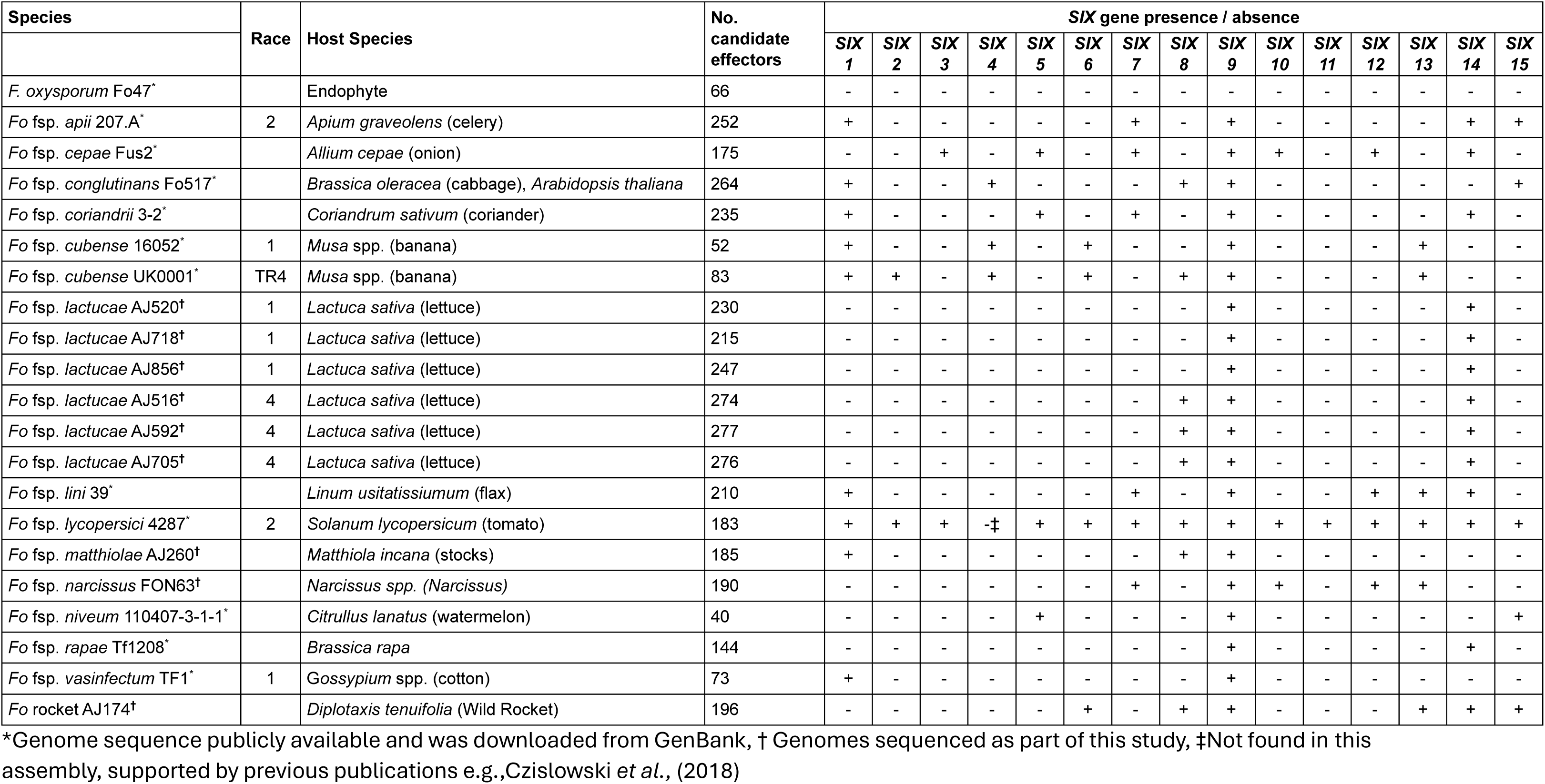
Effector candidates and *SIX* gene distribution for *Fusarium oxysporum* genomes. Candidate effectors identified using TBLASTN (1e-6 cut-off, 65% identity and 65% coverage) of *Fusarium* isolate *mimp*-associated ‘pan-effectorome’ followed by SignalP and EffectorP filtering. *SIX* genes identified using TBLASTN (cut off 1-e^6) using *SIX* genes from *F. oxysporum* f.sp. *lycopersici* isolate 4287 as a reference.

Candidate effectors were clustered into groups using CD-HIT (65% identity) resulting in 238 clusters, including 101 candidate effector clusters for Fola. Of these, 48 were shared amongst all Fola isolates, 18 were unique to Fola4 while 11 were unique to Fola1. The 24 remaining Fola candidate effector clusters (CECs) did not show a consistent presence based on race (Figure 2). Of these 24 non-race specific clusters, eight displayed an interesting distribution pattern within the Fola4 isolates. Candidate effector clusters CEC_153, CEC_173, and CEC_232 were found in Fola4 AJ516, but not in Fola4 AJ592 or AJ705, whereas candidate effector clusters CEC_10, CEC_32, CEC_185, CEC_212 were present in Fola4 AJ592 and AJ705 but not identified in Fola4 isolate AJ516. Of these clusters, CEC_10, CEC_212, and CEC_232 were found in all of the Fola1 isolates, while the remaining clusters (CEC_32, CEC_153, CEC_173, 185) were not identified in any of the Fola1 isolates. Effector complement therefore distinguished between Fola1 and Fola4 isolates, but also suggested that there were two variants of Fola4, one represented by isolate AJ516 and the other by isolates AJ705 and AJ592.

**Figure 2.**
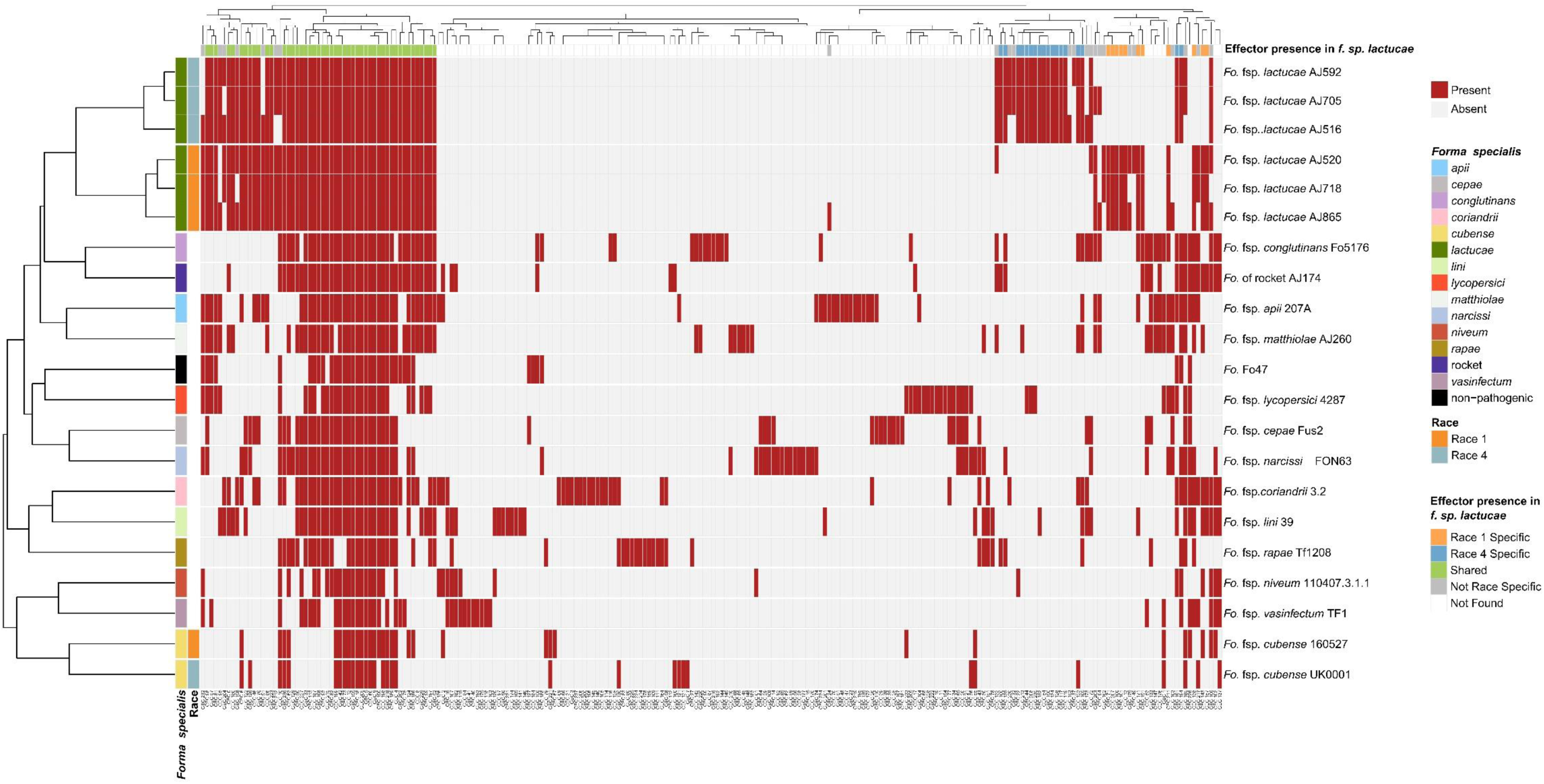
*Fusarium oxysporum* f.sp. *lactucae* races 1 and 4 and other *F. oxysporum* f.spp. clustered based on *mimp*-based effector profile. *Mimps* were identified in the *F. oxysporum* assemblies using either a custom Python script or NHMMER (version 3.3.1) search using a *mimp*-profile Hidden Markov Model. Predicated gene models (Augustus, version 3.3.3) and open reading frames (EMBOSS getorf, version 6.6.6) were filtered for a predicted signal peptide using SignalP (version 4.1) and clustered (CD-HIT, version 4.8.1) to generate a non-redundant protein set for each assembly. Likely effectors were identified using EffectorP (version 2.0.1) and searched against the assemblies using TBLASTN (1e-6 cut-off, 65% identity and 65% coverage). Blast hits were then extracted, sequences filtered (SignalP and EffectorP, default parameters), and clustered into effector groups at ≥65% identity), generating a binary data matrix used to create a heatmap (R Package, Pheatmap). Red indicates effector presence, grey indicates absence.

### *SIX* gene phylogenies reveal variation between Fola1 and Fola4, and differences in *SIX8* between Fola4 isolates

Fola races displayed different *SIX* gene profiles with homologs of *SIX8, SIX9,* and *SIX14* in all Fola4 isolates, and homologs of *SIX9* and *SIX14* in all Fola1 isolates. Phylogenetic analysis of *SIX8*, *SIX9*, and *SIX14* revealed sequence variation between Fola races in *SIX9* and *SIX14* as well as variation in *SIX8* sequences amongst the Fola4 isolates (Figure 3). Fola4 isolates had a single copy of *SIX8* which was identical for isolates AJ705 and AJ592, but different for isolate AJ516, with 16 base substitutions and one indel from position 528 to 535 (supplementary data SD1-T4). The Fola4 *SIX8* sequences were most closely related to the *F. oxysporum* isolate from rocket (AJ174), *F. oxysporum* f. sp. *conglutinans* (Fo5176), and *F. oxysporum* f. sp. *matthiolae* (AJ260 and PHW726 1). The two different *SIX8* sequence variants within Fola4 were consistently identified within 29 isolates obtained from Italy, Ireland, Netherlands, Spain and UK following PCR and sequencing with 23 isolates having the same sequence as Fola4 isolate AJ516 and 6 isolates having the AJ592 / AJ705 sequence (data not shown). Two copies of *SIX9 (SIX9.1 and SIX9.4)* were identified in the Fola1 isolates, with four copies identified in Fola4 isolate AJ516, and five copies in Fola4 isolates AJ705 and AJ592. There was variation in the two copies of *SIX9* in the Fola1 isolates, which were in separate clades, but there was no variation in these copies between isolates (Figure 4). One of the copies of *SIX9 (SIX9.4)* appears to have been duplicated in Fola4, with two copies of *SIX9.4* in each isolate identical to the Fola1 copy. The two additional copies of *SIX9* (*SIX9.2 and SIX9.3*) found in all Fola4 isolates but were not identified in Fola1 and were similar to copies of *SIX9* identified in *F. oxysporum* from rocket (AJ174), *F. oxysporum* f. sp. *conglutinans* (Fo5176), and *F. oxysporum* f. sp. *matthiolae* (AJ260 and PHW726 1), as observed for *SIX8.* Copies of *SIX9* from *F. oxysporum* f.spp. *luffae, coriandrii,* and *capsici* were also similar to *SIX9.2 and SIX9.3* identified in Fola4. The remaining sequence variant of *SIX9* (*SIX9.1*) identified was only present in Fola4 isolates AJ705 and AJ592 but was absent in Fola4 isolate AJ516, where the different *SIX8* sequence was identified in comparison with Fola4 isolates AJ705 and AJ592. All Fola4 isolates each had one identical copy of *SIX14* which was the same as the two copies of *SIX14* present in each of the Fola1 isolates (Figure 5). The Fola1 isolates also had an additional third copy of *SIX14.* The different sequence variants of *SIX14* in all Fola isolates were more closely related to each other than to homologues of *SIX14* in other *F. oxysporum* f.spp.

**Figure 3:**
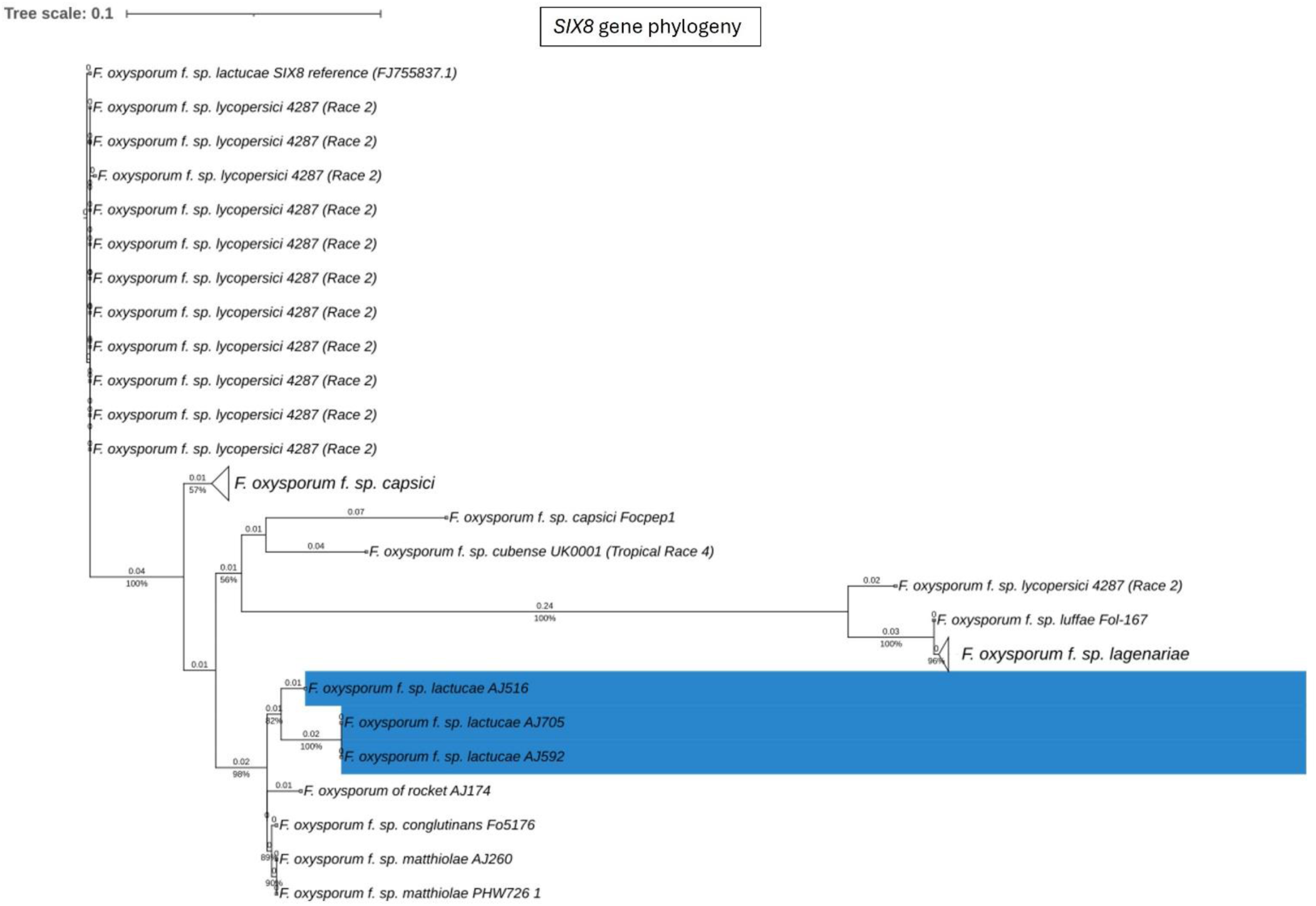
Phylogenetic tree of *Fusarium oxysporum SIX8* sequences. ModelFinder, using Bayesian information criterion, was used to calculate the best model of sequence evolution, selecting K2P+G4. IQ-tree2 was used to estimate the maximum likelihood (ML) tree under this model, with 1000 ML bootstrap replicates expressed as a percentage (UFBoot2). The tree is rooted through the *F. oxysporum* f.sp. *lycopersici* SIX8 reference FJ755837.1. *F. oxysporum* f.sp. *lactucae* race 4 isolates are highlighted in blue. Branch lengths are shown above the branch and the scale bar indicates 0.1 substitutions per site. Collapsed clades are denoted by the white triangle.

**Figure 4:**
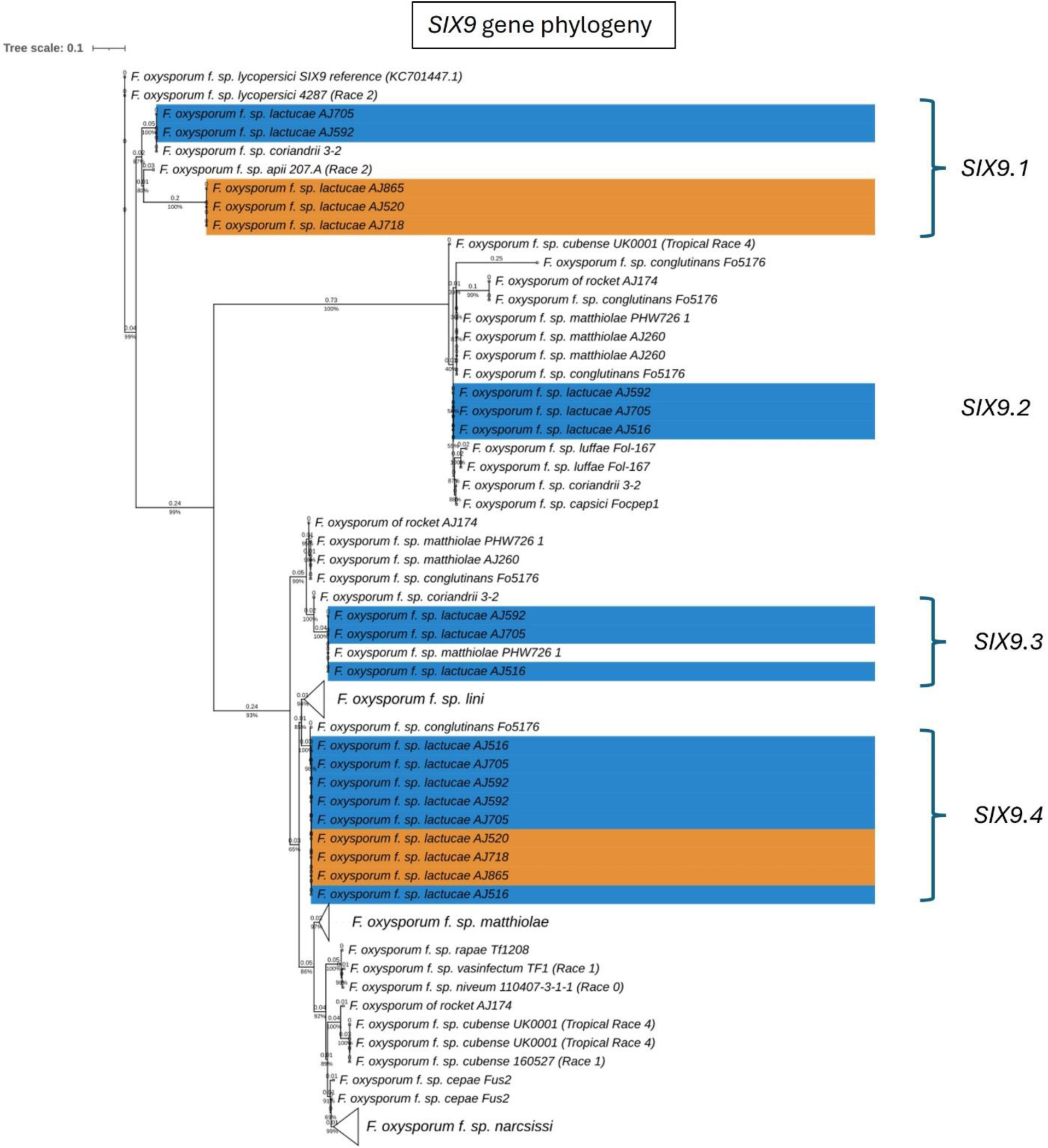
Phylogenetic tree of *Fusarium oxysporum SIX9* sequences. ModelFinder, using Bayesian information criterion, was used to calculate the best model of sequence evolution, selecting TPM3+G4. IQ-tree2 was used to estimate the maximum likelihood (ML) tree under this model, with 1000 ML bootstrap replicates expressed as a percentage (UFBoot2). The tree is rooted through the *F. oxysporum* f.sp*. lycopersici SIX9* reference KC701447.1. *F. oxysporum* f.sp. *lactucae* race 1 and race 4 isolates are highlighted in orange and blue respectively. Branch lengths are shown above the branch and the scale bar indicates 0.1 substitutions per site. Collapsed clades are denoted by the white triangle.

**Figure 5:**
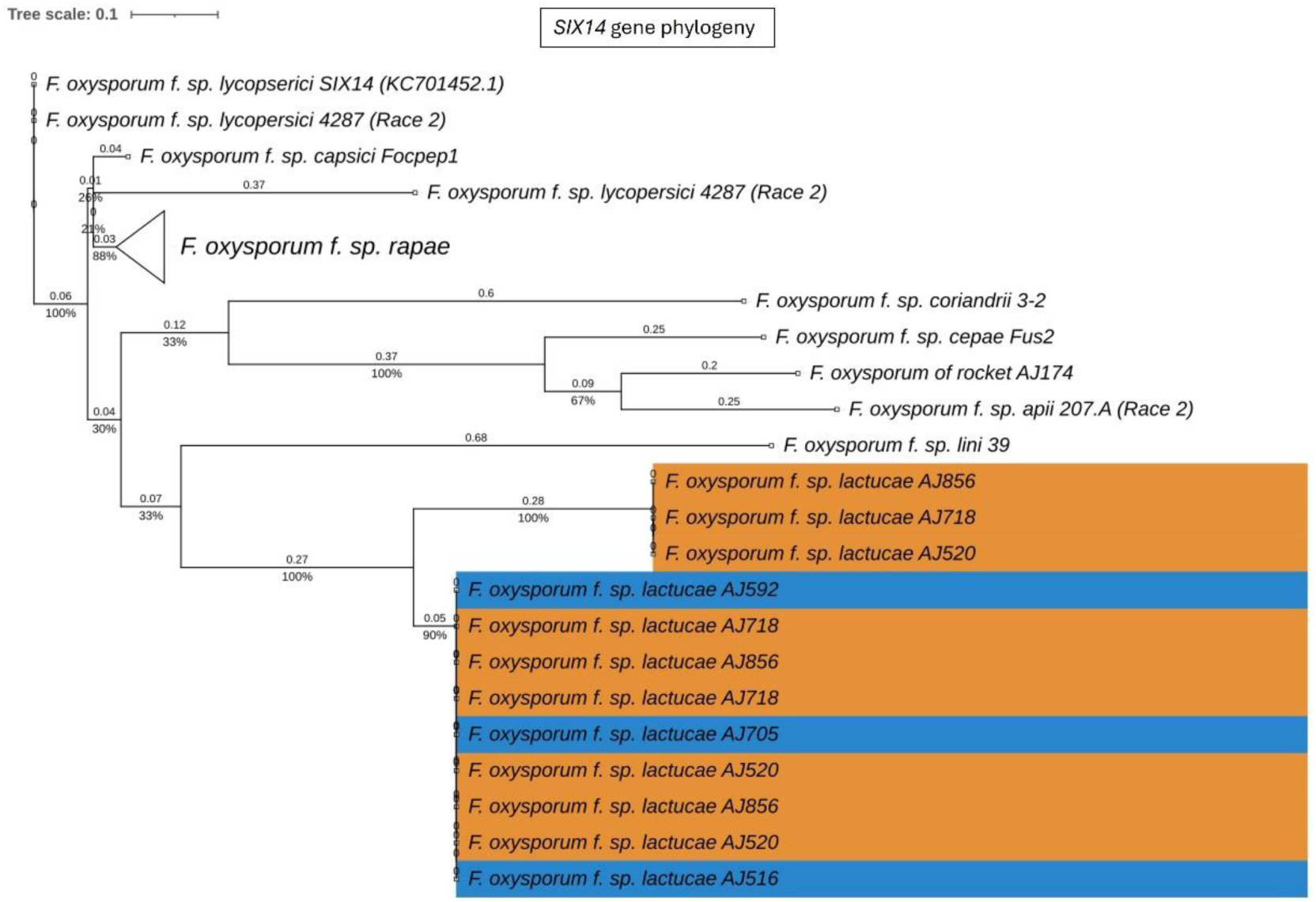
Phylogenetic tree of *Fusarium oxysporum SIX14* sequences. ModelFinder, using Bayesian information criterion, was used to calculate the best model of sequence evolution, selecting K2P+G4. IQ-tree2 was used to estimate the maximum likelihood (ML) tree under this model, with 1000 ML bootstrap replicates expressed as a percentage (UFBoot2). The tree is rooted through the *F. oxysporum* f.sp. *lycopersici SIX14* reference KC701452.1. *F. oxysporum* f.sp. *lactucae* race 1 and race 4 isolates are highlighted in orange and blue respectively. Branch lengths are shown above the branch and the scale bar indicates 0.1 substitutions per site. Collapsed clades are denoted by the white triangle.

In summary, the *SIX* gene phylogenies showed clear differences between the two Fola races in terms of gene copy number and sequence; the absence of *SIX8* in Fola1, only two *SIX9* gene variants in Fola1 compared to up to four *SIX9* gene variants in Fola4 isolates, and three *SIX14* gene copies in Fola1 compared to a single copy of *SIX14* in Fola4. Furthermore, differences between Fola4 AJ516 *SIX8* gene sequence and *SIX9* gene variants, compared to AJ705 and AJ592, supported the evidence from the genome phylogenies and the candidate effector profiles for two variants of Fola4.

### Collinearity analysis highlights differences in accessory genomes of Fola isolates

To further understand genome evolution across the Fola isolates, we determined large scale synteny across both core and accessory regions through collinearity analysis. When Fola core sequences were aligned to the Fol 4287 core chromosomes, all Fola isolates demonstrated high collinearity (Figure S1), as has been previously observed in other *F. oxysporum* f.spp.

However, we observed a possible translocation event in Fola1 AJ520, whereby sequence syntenic with Fol 4287 chromosome 9 was joined to AJ520 contig 2, the rest of which was syntenic with Fol 4287 chromosome 5. This potential translocation region was supported by approximately 30x coverage of long read sequencing data (data not shown) but was not investigated any further. When Fola accessory genome sequences were examined, all three of the Fola1 isolates were found to share a high level of collinearity, with large blocks of syntenic sequences and evidence of some structural variations across the larger contigs (Figure 6). However, collinearity analysis between Fola1 AJ520 and Fola4 AJ516 demonstrated an overall absence of large-scale synteny between these two races, with only small-scale alignments scattered throughout both isolates (Figure 6). Fola4 AJ592 and AJ705 also demonstrated high collinearity, with the majority of contigs being highly syntenic and evidence of only a small number of rearrangements. However, the relationship between Fola4 AJ705 and AJ516 was much more variable. While some contigs shared large blocks of synteny, the remaining contigs displayed evidence of numerous structural variations, including large fragment inversions, translocations, insertions and deletions. These results further highlighted that despite the high collinearity across the core genome sequences, much more genomic variation was observed throughout the accessory genome sequences across the Fola isolates with clear lack of synteny between Fola1 and Fola4. The results also again identified differences between the two Fola4 variants represented by AJ516 and AJ705 / AJ592.

**Figure 6.**
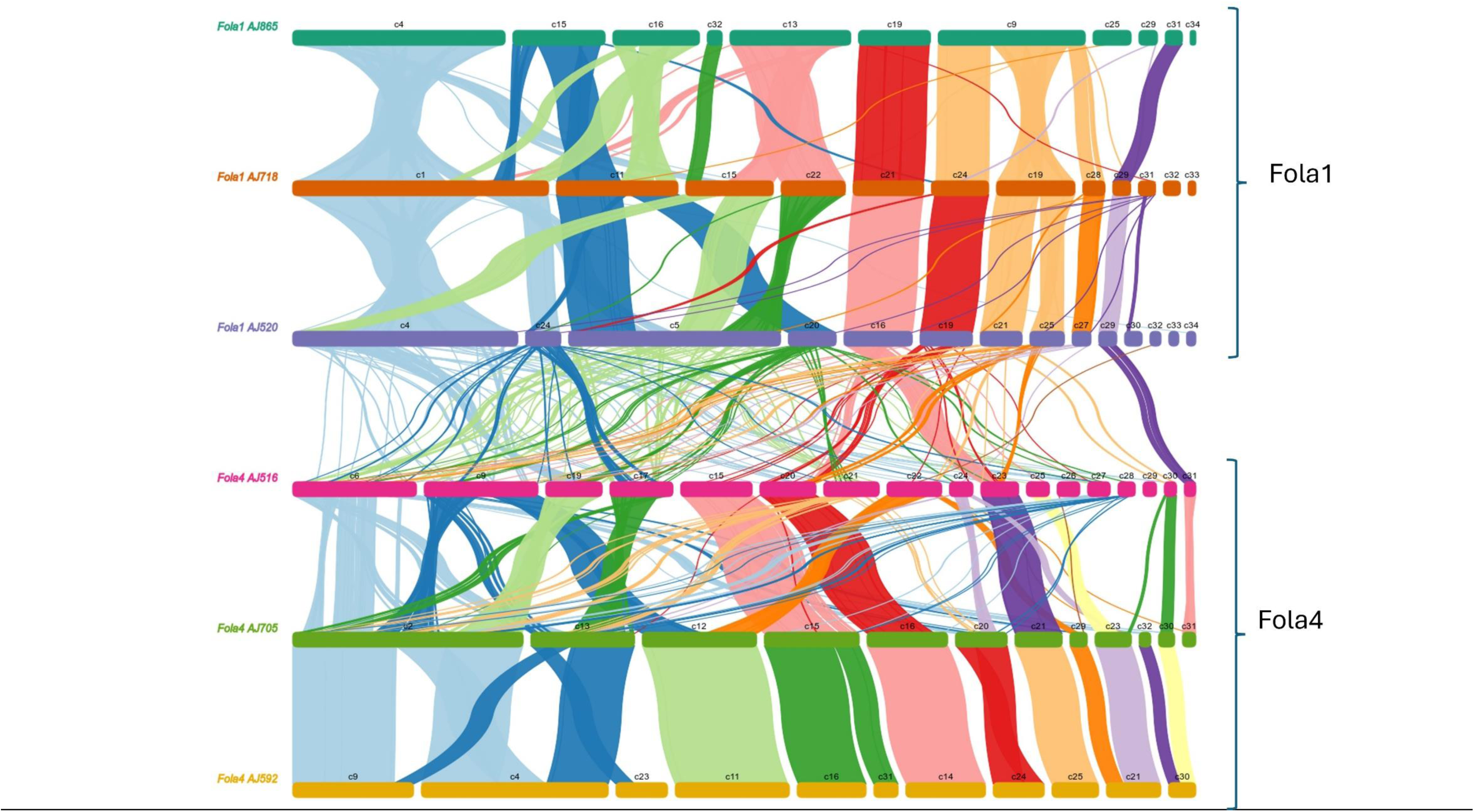
Collinearity analysis of *Fusarium oxysporum* f.sp. *lactucae* (Fola) race 1 and race 4 accessory genome sequences, showing syntenic alignments larger than 10 Kb.

### Fola accessory chromosomes are enriched for *mimps*, predicted effectors, *SIX* genes and transposable elements

Genome organisation in Fola1 and Fola4 followed a similar pattern to previously reported *F. oxysporum* genomes (Armitage et al., 2018). *Mimp* sequences (Figure 7, track B), which have previously been shown to be associated with known effectors in *F. oxysporum* f.spp. (Schmidt et al., 2013), were enriched on accessory contigs, notably contigs 9, 6, 15, 17, 19, 20, 22 and 27 of Fola4 AJ516, and contigs 4, 5, 19, 24, 27 and 29 of Fola1 AJ520 (Figure 7, track B).

**Figure 7.**
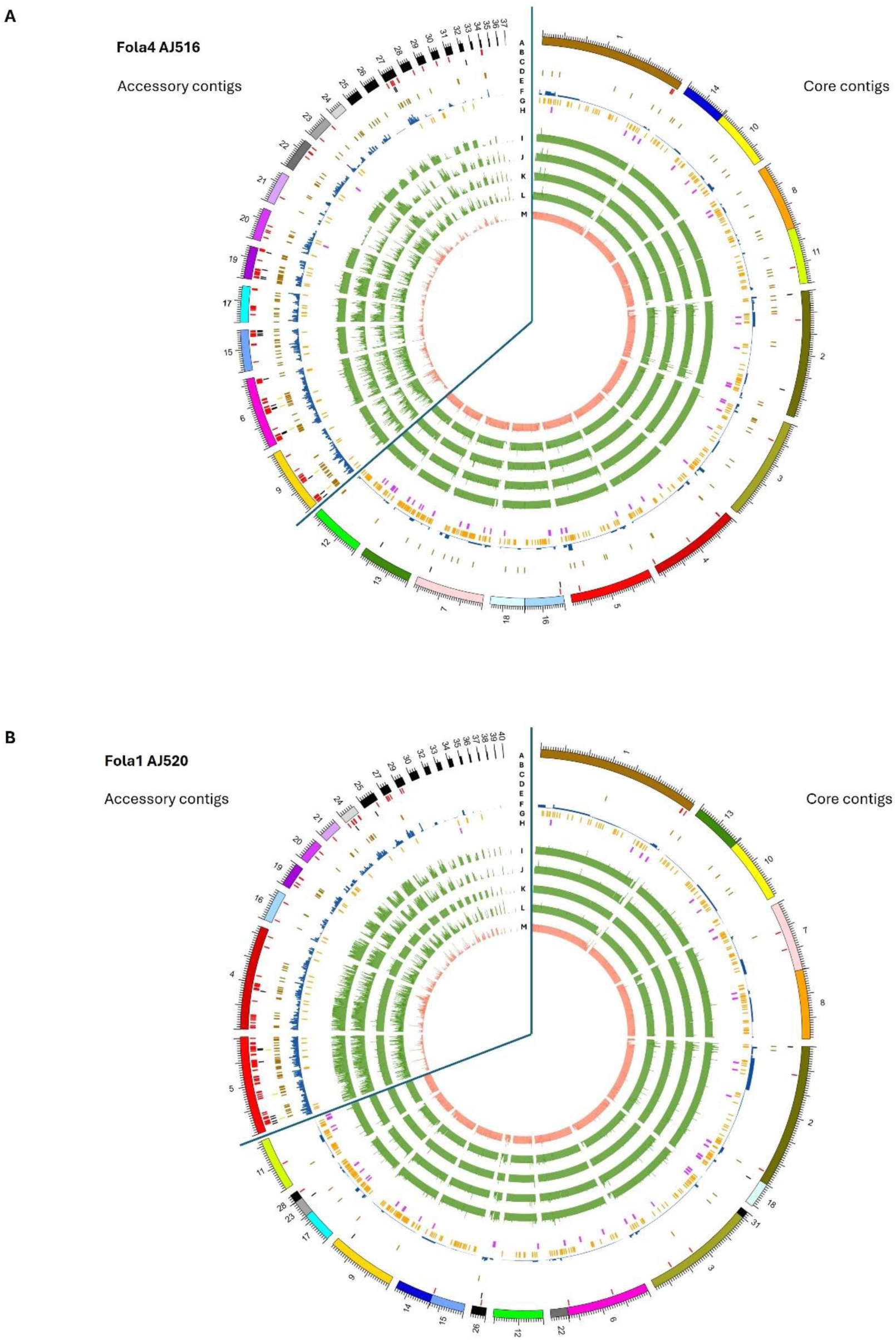
Genomic features on core and accessory contigs for (A) *Fusarium oxysporum* f. sp. *lactucae* race 4 (Fola4) isolate AJ516 and (B) *F. oxysporum* f. sp. *lactucae* race 1 (Fola1) isolate AJ520. Inner tracks moving in from the outside (Track A) genome assembly contigs, (Track B) *mimp* sequences, (Track C) predicted effector genes (Track D) *SIX* gene homologues (Track E) Helitron transposable elements f) other transposable elements (Track F) secreted carbohydrate active enzymes (Track H) secondary metabolite clusters. Inner tracks (I-M) showing alignment of assemblies to **A)** Fola4 AJ516 from Fola4 AJ705 (Track I), Fola1 AJ520 (Track J), Fola1 AJ865 (Track K), Foma AJ260 (Track L) and non-pathogen Fo47 (Track M), and **B)** Fola1 AJ520 from Fola4 AJ516 (Track I) Fola4 AJ705 (Track J), Fola1 AJ865 (Track K) Foma AJ260 (Track L) and non-pathogen Fo47 (Track M).

*SIX* gene homologues were located on contigs 6 and 9 for Fola4 AJ516 (Figure 7A, track D) and contig 5 for Fola1 AJ520 (Figure 7B, track D) indicating that these are putative pathogenicity contigs. Predicted *mimp*-associated effector genes were found in clusters on contigs 6, 9, 15, 19 and 27 for Fola4 AJ516, and on contig 5 in Fola1 AJ520 (Figure 7, track C). Helitrons and other transposons were also enriched on accessory contigs (Figure 7, tracks E, F). In contrast, the frequency of secreted carbohydrate active enzymes (CAZymes) and secondary metabolite clusters (track H) were reduced on the accessory contigs compared to the core (Figure 7, track G). Fola4 AJ516 core contigs 7, 13 and 12 and Fola1 AJ520 core contigs 9, 11 (and 17, 23, 28) showed enrichment of CAZymes as observed for *F. oxysporum* f.sp. *cepae* by Armitage et al. (2018). Very few *mimps* or predicted effectors were present on Fola core contigs. Although helitrons were enriched on accessory contigs, they were also present on core contigs for both Fola1 AJ520 and Fola4 AJ516. Core contigs showed a consistent level of alignment across all comparative genomes (Fola4 AJ516 and AJ705, Fola1 AJ520 and AJ865, Foma AJ260 and Fo47; Figure 7, tracks I - M). In contrast, accessory contigs of Fola4 AJ516 and Fola1 AJ520 generally showed the most consistent levels of alignment with isolates of the same race, while more variable alignment indicating duplication or loss events, was evident with isolates of the opposite Fola race (Figure 7A tracks J-K, 7B tracks I-J) or with Foma AJ260 (track L). Low levels of alignment were observed for all Fola isolates with the non-pathogen Fo47 (Figure 7, track M). Interestingly, accessory contigs 21 and 28 from Fola4 AJ516 (Figure 7A) show a lower level of alignment against Fola4 isolate AJ705 (track I) than it does to either Fola1 AJ520 (track J) or Foma AJ260 (track L), with levels of alignment against Fola4 AJ705 being more similar to the non-pathogen Fo47 for these two contigs. On further investigation it was found that the subtelomeric 7.5 kb sequences of these two contigs (5’ telomere for contig 21 and 3’ telomere for contig 28) differed to all other subtelomeric sequences of AJ516, but were very similar to each other. This suggests that these two contigs may be related and possibly part of the same chromosome. Subtelomeric sequences are rapidly evolving regions of the genome (Rahnama, et al., 2021) which tend to differ between isolates of different *F. oxysporum* f.sp. while being very similar within a f.sp. (Huang, 2023). Phylogenetic analysis of all sub-telomere 7.5 kb regions found in Fola4 AJ516, AJ705 and Fola1 AJ520, showed that Fola1 subtelomeric sequences clustered separately from Fola4 isolates, while Fola4 isolates AJ516 and AJ705 subtelomeric sequences clustered together with the notable exception of AJ516 contigs 21 and 28 which clustered separately (Supplementary Figure S2). This suggests that contigs 21 and 28 from Fola4 AJ516 may have been acquired by HCT from an unknown donor. Although we do not know how recently this HCT event occurred, this provides further evidence that separates the two Fola4 variants.

### RNAseq identifies key shared and unique effectors expressed by Fola1 and Fola4 in lettuce

RNAseq data for Fola1 AJ520 and the isolates representing the two variants of Fola4 (AJ516 and AJ705) were filtered to identify differences in putative effectors differentially expressed *in planta* (supplementary data SD1-T6-T11, Figure 8). This analysis identified 14 putative expressed effectors shared between Fola1 AJ520 and Fola4 AJ516 and AJ705, including the known effectors *SIX9* and *SIX14* (Figure 8). Of these effectors, all but one (OG0017028) showed homology to hypothetical proteins or putative effector proteins from other members of the FOSC following a blastx search of the NCBI database, with several homologues identified in *F. oxysporum* f.spp. *apii*, *raphani*, *cubense* and *narcissi*. Seven putative effectors were identified as Fola4 specific, including the previously reported divergently transcribed effector pair *SIX8* and *PSE/PSL1* (Ayukawa et al., 2021), and one of the *SIX9* homologues.

**Figure 8.**
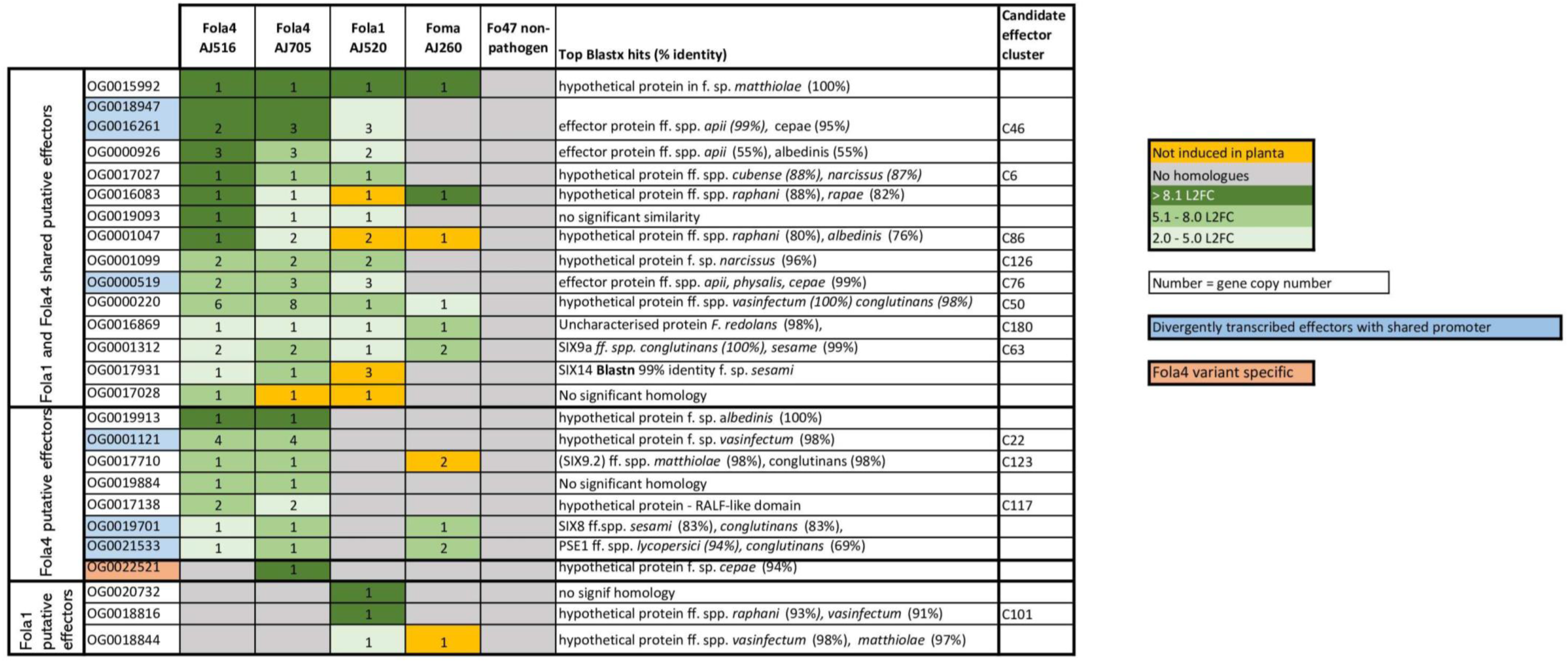
Filtered shared and unique candidate effectors upregulated *in planta* for *Fusarium oxysporum* f.sp. *lactucae* race 1 (Fola1) and race 4 (Fola4) isolates in comparison with *Fusarium oxysporum* f.sp. *matthiolae* (Foma) and non-pathogenic *F. oxysporum* Fo47. Candidate effectors are listed by orthogroup and matched to the relevant candidate effector cluster on the effector heatmap (Figure 2) where applicable. Orthogroup numbers coloured blue indicate genes with divergently transcribed paired effectors. Orthogroup OG0022521 (highlighted in pink) is only present in Fola4 AJ705 and AJ592. Numbers indicate gene copy number. Colours indicate level of induction by log 2 fold change (L2FC). Dark green = L2FC >8.1, mid-green = L2FC 5.1 - 8.0, light-green = L2FC 2.0 - 5.0, orange = no induction *in planta*, grey = no homologues.

An additional effector (OG0022521) was identified as present and expressed only in Fola4 AJ705 with homology to a hypothetical protein in *F. oxysporum* f.sp. *cepae*. Three putative effectors were identified as specific to Fola1, two of which had homologues in *F. oxysporum* f.spp. *vasinfectum*, *raphani* and *matthiolae*.

*SIX8, SIX9* and *SIX14* genes were located on contigs 6 and 9 of Fola4 AJ516, with the other expressed putative effectors located mainly on contigs 6, 9, with a few on 15, 19, 27 and 34. In Fola4 AJ705, *SIX* genes were located on contig 2 and contig 20, with the majority of expressed putative effectors located on contig 2, with fewer on contigs 20, 24, 15, 12, and 13. *SIX9* and *SIX14* in Fola1 AJ520 were located on contig 5, with other expressed putative effectors also located mainly on contig 5, with some on contig 4. This identified the main putative pathogenicity contigs as contigs 6 and 9 in Fola4 AJ516, contigs 2 and 20 in Fola4 AJ705 (corresponding to contigs 9, 4 and 24 in Fola4 AJ592) and contigs 4 and 5 for Fola1 AJ520.

Several of the highly expressed putative effectors were found to be arranged as gene pairs divergently transcribed from a shared promoter. This included the Fola4 specific genes *SIX8* and *PSL1*, and the gene pair OG0000519 - OG0018947/OG0016261, which showed evidence of duplication, being present in Fola1 AJ520 and Fola4 AJ705 as three copies and in Fola4 AJ516 as two copies. In Fol, *Avr2* (*SIX3*) is also arranged as a paired effector with *SIX5* sharing a promoter (Cao et al., 2018). Both *SIX8* and *Avr2* are members of the same ToxA-like structural family (Yu et al., 2024) with their paired partner (*PSE1*/*PSL1* and *SIX5* respectively) being members of another structural group, family 4. Further analysis will show if the newly identified paired putative effectors present in Fola will fall into the same categories. In addition, Fola4 carried a head-to-head pair of identical genes with a shared promoter that is duplicated to give four copies of the gene (OG0001121). Moreover, orthogroups OG0000926, OG0001047, OG0001099, OG0000220, OG0001312 and OG0017710 were all present as multiple copies showing evidence of duplications.

Orthogroup analysis also highlighted multiple differences in copy number and sequence variation for some of the expressed putative effectors. Of the multiple *SIX9* copies identified in the DNA phylogeny in the two Fola races (Figure 4), only *SIX9.2* (present in Fola4 only) and *SIX9.4* (present in all Fola isolates) were differentially expressed *in planta.* For the *SIX9.1* copy (present in Fola1 and Fola4 AJ592 / AJ705 only), it was found that although the full gene was annotated and present in Fola4 AJ705 and AJ592, it was not expressed *in planta*. Only a partial copy of SIX9.1 was present on contig 5 in Fola1 (which was not annotated) with no RNAseq reads aligned to it. The Fola4 *SIX9.3* gene copies were all partial copies with no gene annotation. In summary, Fola1 AJ520 contained one full-length *SIX9* gene copy (*SIX9.4*) that is upregulated *in planta* and identical to a duplicated upregulated *SIX9* gene copy in Fola4 isolates (*SIX9.4*). Fola4 isolates also contained an additional *SIX9* gene copy (*SIX9.2*) that was also upregulated *in planta*.

Although *SIX14* (OG0017931) had identical DNA sequences for both Fola races, there were differences in both copy number and expression level between Fola1 and Fola4. Despite Fola1 AJ520 having three copies of *SIX14* all located on accessory contig 5, RNAseq indicated that there was low basal expression of these genes with only 30 transcripts per million (TPM) and no upregulation *in planta* (10 TPM). In contrast, the single copy of *SIX14* found in Fola4 AJ516 (contig 9) showed a L2FC of 4.3 with high level of expression, rising from a basal level of 483 TPM to 6309 TPM *in planta* (Supplementary data SD1). Further investigation of the genome showed that Fola4 AJ516 *SIX14* has a *mimp* sequence 151 bp upstream of the start codon, while the two full-length copies of *SIX14* in Fola1 AJ520 on contig 5 have a helitron transposon inserted into the promoter region immediately upstream of the start codon. The third copy of *SIX14* in Fola1 AJ520 has transposon Tf2-9 inserted into the intron between exon 1 and exon 2. In addition, the *mimp* sequence is 1 kb upstream of the start codon. Thus, all copies of *SIX14* in Fola 1 have disruptions or changes to the promoter region with one copy also having a disruption to the gene sequence which prevents expression of this putative effector *in planta*. Other orthogroups that showed differences in expression between races (OG0016083, OG0001047, and OG0019028) require further investigation.

### Fola4 isolates show differences in SIX8-PSE1/PSL1 effector pair sequence

The putative effector gene *PSE1* has previously been reported as divergently transcribed *in planta* from the same promoter as *SIX8* in *F. oxysporum* f.sp. *conglutinans* (Focn, isolate Cong1-1), with both genes being required for virulence on *Arabidopsis* (Ayukawa 2021). Furthermore, the authors reported a similar gene, *PSL1* (PSE1-like) paired with *SIX8* in the tomato infecting isolate Fol 4287 that differed in ten amino acids at the C-terminal (Ayukawa, 2021). Both *SIX8* and *PSE1* / *PSL1* divergently transcribed genes were identified as putative effectors that were induced *in planta* for both Fola4 isolates AJ705 and AJ516 in our RNAseq analysis. As described previously, Fola4 isolates all contain homologues of *SIX8* that are closely related to those present in Focn, Foma and *F. oxysporum* from rocket. These *SIX8* sequence differences identified between Fola4 AJ516 compared with AJ705 and AJ592 result in 7 amino acid substitutions in the SIX8 protein sequence (Figure 9A). All Fola4 isolates also showed differences in the amino acid sequence of *SIX8* compared to Focn and Foma, with Fola4 AJ516 having 4 amino acid substitutions and AJ705 / AJ592 having 3 different amino acid substitutions. Alignment of the PSE1 and PSL1 protein sequences (Figure 9B) showed that the PSL1 found in Fola4 AJ516 is more closely related to that from the tomato infecting isolate Fol 4287 than to the PSE1 in either of the other Fola4 isolates AJ705 or AJ592 which in turn are identical to the PSE1 from Focn isolate 5176 and the Foma AJ260 isolate. Notably, isolate Fola4 AJ516 shares the same ten amino acid difference at the C-terminus of PSL1 as the tomato infecting isolate Fol 4287 as previously described (Ayukawa, 2021). DNA sequence alignment of a 4 kb region encompassing the *SIX8-PSL1* gene pair (Figure 9C) from all Fola4 isolates, Fol 4287, Focn 5176 (100% sequence identity to isolate Cong1-1 for the *SIX8 - PSE1* gene pair locus), Foma AJ260 and *F. oxysporum* AJ174 from rocket showed that sequence homology between these *F. oxysporum* f.spp. is confined to the region of the genes and, to a lesser extent, the intergenic region, while alignment was quickly lost outside of this region. Interestingly, the only *F. oxysporum* isolate that had a full *mimp* sequence present in the shared promoter was Foma AJ260 (Figure 9C). All isolates that harboured the *SIX8-PSE1/PSL1* gene pair, had a *mimp*TIR. Furthermore, sequence alignment between the Fola4 isolate AJ516 compared to isolates AJ705 and AJ592 was also lost outside of the *SIX8-PSL1* gene pair. The *SIX8-PSE1* gene pair is located on contig 20 of AJ705 and contig 24 of AJ592 which are both telomere to telomere contigs with 99.92% identity to one another. These 1.12 Mb chromosomes also harboured the *SIX9.1* gene copy that is absent in AJ516. Overall these data strongly suggest that Fola4 AJ705 has potentially gained the *SIX8-PSE1* gene pair (along with *SIX9*.1 gene copy) by HCT or HGT from a different source to that for the Fola4 AJ516 *SIX8-PSL1* gene pair. This is again supporting evidence for two variants of Fola4. In addition, collinearity analysis (Figure 6) identified a region of synteny between Fola4 AJ516 contig 6 and Fola4 AJ705 contig 9 that appears to span the location of AJ516 *SIX8*-*PSL1*. Further investigation found that 39 kb of additional genetic material was present in Fola4 AJ516 at the site of a *F. oxysporum* specific helitron sequence in Fola4 AJ705 (Figure 9D). The additional 39 kb present in Fola4 AJ516 was flanked by identical helitrons to that present at this site in AJ705. These helitrons have similar terminal sequences to FoHeli1 (Chellapan et al., 2016) and contain intact rep and hel domains for autonomous replication. The arrangement of the FoHeli sequences and insertion sites suggests either gene gain in Fola4 AJ516 or gene loss in Fola4 AJ705, possibly by homologous recombination at the helitron site (Figure 9D and E). Helitrons usually insert at a TA site so the immediate 5’ nucleotide to the helitron sequence is usually a T and 3’ is usually an A. This was not the case for Fola4 AJ705 which has a TT insertion site, with AAT-5’ and 3’-TGT flanking FoHeli1a (Figure 9E). In Fola4 AJ516 the FoHeli1b 5’ matches the AA T-5’ of AJ705 FoHeli1a and the FoHeli1c 3’-TGT matches AJ705 FoHeli1a. Recombination between FoHeli1b and FoHeli1c could therefore have produced a single FoHeli1 as in AJ705 FoHeli1a, thus losing the *SIX8-PSL1* gene pair.

**Figure 9.**
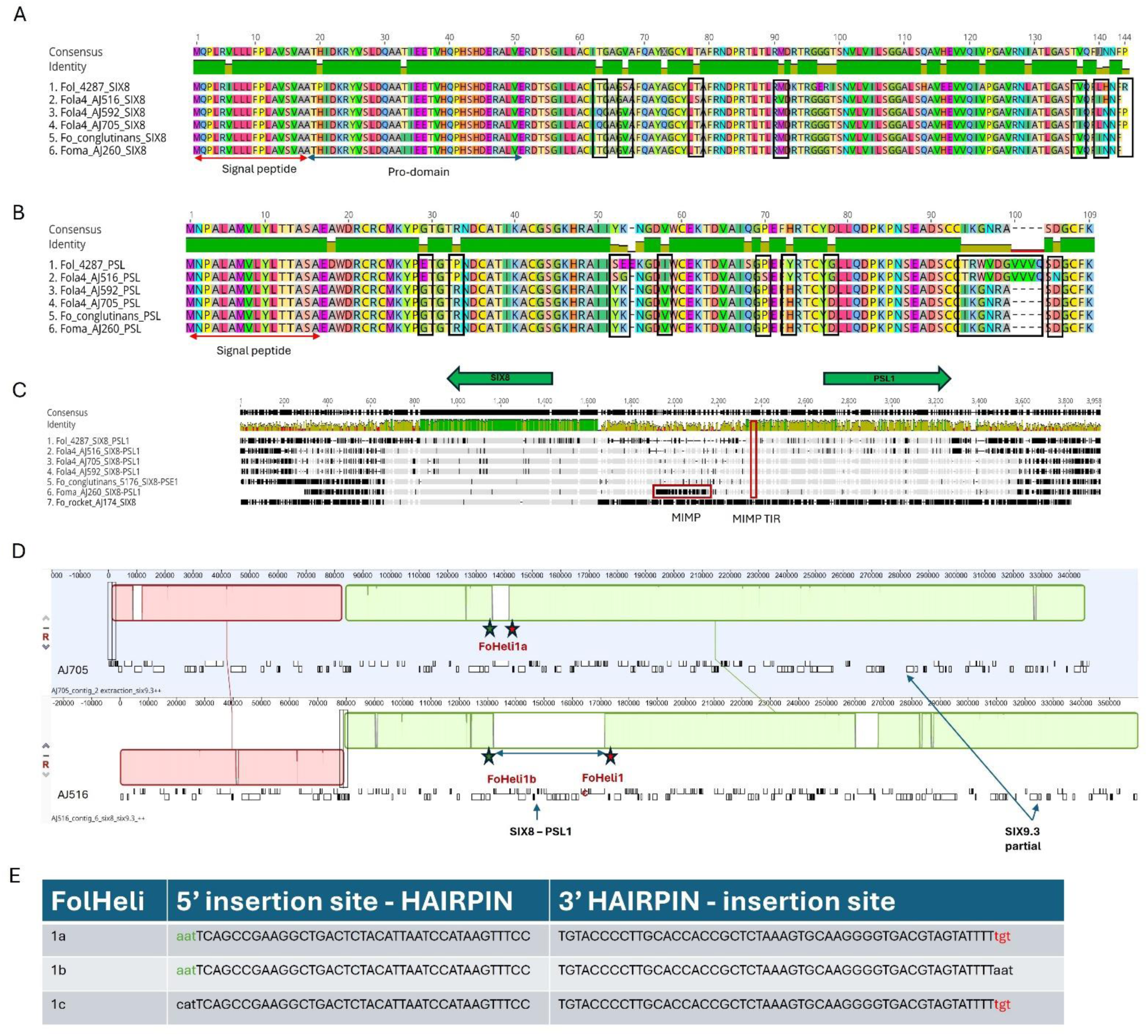
Comparison of *SIX8* and *PSE1/PSL1* gene pair partition for two *Fusarium oxysporum* f.sp. *lactucae* race 4 (Fola 4) variants represented by isolates AJ516 and AJ592 / AJ705 compared with *F. oxysporum* f.sp. *conglutinans* (Focn) isolate 5176 and *F. oxysporum* f.sp. *mathiolae* isolate (Foma) isolate AJ260. Geneious Prime (2023.2.1) alignment of **A)** SIX8 and **B)** PSE1/PSL1 protein sequences for Fol 4287, Fola4 AJ516, AJ705 and AJ592, Focn 5176 and Foma AJ260. Amino acid differences between the two Fola4 subgroups are outlined in black. **C)** Geneious alignment using progressive pairwise alignment and the neighbour-joining method of tree building on default settings to create the tree was carried on the 4kb genomic region covering the *six8 – psl1* genes*. mimp* and *mimp*TIR sequences are outlined in red. **D)** Anchored genome alignment of locally collinear blocks of 350 kb genomic regions for Fola4 AJ705 contig 2 using Mauve default settings within Geneious Prime 2023.2.1 (top panel) and AJ516 contig 6 (bottom panel). The location of identical FoHeli1-like helitrons are shown. The green and red stars show location of identical 5’ and 3’ insertion sequences respectively. **E)** 5’ and 3’ insertion sequences of each FoHeli1 with FoHeli1 termini shown in capitals and genomic sequence in lower case.

## Discussion

### Fola1 and Fola4 have emerged independently from a common ancestor

This research has presented the first full genome and transcriptome data for Fola, which has enabled the identification of key differences in effector repertoire and expression in the globally significant races Fola1 and Fola4 for the first time. Four lines of evidence suggest that the two races emerged independently (from a common ancestor) rather than the newly identified Fola4 evolving from Fola 1 for instance. Firstly, phylogenetic analysis of both core and accessory genomes indicated that Fola1 and Fola4 were in sister clades derived from a common ancestor. This is also supported by a previous report where Fola1 and Fola4 isolates were consistently shown to be in distinct vegetative compatibility groups (Pintore et al., 2017) and by GBS based phylogenetic analysis (Claerbout et al., 2023). Secondly, the *mimp*-associated effector profile for Fola1 and Fola4 also separates them into separate clusters.

However, of those predicted effectors that are shared between the two races, the majority are also present in the non-pathogen Fo47 along with many other f.spp. and so are not pathogen or host specific. A third line of evidence which supports the independent emergence of Fola4 is the collinearity analysis of the accessory genome. Here, the larger accessory genome size of Fola4 (17.7 – 20.9 Mb) compared with Fola1 (15.2 – 15.5 Mb) might suggest that Fola4 arose from Fola1 by gain of additional genetic material via HCT / HGT; however, collinearity analysis showed large blocks of synteny between isolates of the same race, in contrast to extremely fragmented and rearranged small islands of synteny between the two races represented by Fola1 AJ520 and Fola4 AJ516. If Fola4 evolved from Fola1, then larger blocks of synteny might be expected between the accessory contigs that originated from Fola1, especially given the recent emergence of Fola4. A final piece of evidence supporting the independent emergence of Fola4, is the difference in the subtelomeric sequences between the two races. Although telomeric regions are highly variable regions prone to transposon insertion, duplications, and rearrangements (Chiara et al., 2005), some fungi, including *Magnaportha oryzae*, *Metarhizium anisopliae* and *Ustilago maydis*, have remarkably stable telomeric regions that also harbour RecQ-like telomere-linked helicase (TLH) genes within ∼10 kb of the telomere (Sanchez-Alonso and Guzman, 1998; Gao et al 2002; Inglis et al., 2005). More recently, the presence of RecQ-like TLH genes associated with stable, telomere-adjacent sequences specific to *F. oxysporum* f.spp. has been reported (Huang, X., 2023; Salimi et al., 2024) where it was suggested that these regions were common between isolates adapted to the same host. In contrast, preliminary comparisons of sequences of the subtelomeric 7.5 kb region from Fola1 AJ520 and AJ718 with Fola4 AJ516 and AJ705, that includes a RecQ-like helicase gene, suggested that these regions were generally in separate race-specific clusters. Overall, these four lines of evidence support an independent evolution of Fola1 and Fola4 from a shared common ancestor.

### Identification of both shared and race-specific *SIX* genes and novel putative effectors upregulated *in planta* differentiates the Fola races

Genomic and transcriptomic analyses of Fola1 and Fola4 isolates identified different *SIX* gene complements and expression for the first time. All Fola1 and Fola4 isolates harboured copies of *SIX9* and *SIX14*. Of the two copies of *SIX9* found in Fola1, only one (*SIX9.4*), was found to be expressed *in planta*. The Fola 4 isolates contained different copies of *SIX9*; *SIX9.2* and *SIX9.4* were both expressed *in planta* in all Fola4 isolates while *SIX9.3* was a partial gene copy (pseudogene) not annotated in the genomes. Fola4 isolates AJ705 and AJ592 also harboured *SIX9.1* (unlike the other FOL4 variant isolate AJ516) but this was not expressed *in planta*. *SIX9* has been shown to be structurally similar to *SIX11* (Yu et al., 2023) and its presence was recently found to be associated with highly virulent *F. oxysporum* f.sp. *vasinfectum* race 4 isolates (Jobe et al., 2024). Three copies of *SIX14* were identified in the genome of Fola1 AJ520, but transcriptomic analysis showed that none of these were expressed *in planta*, due to disruptions in the promoter and gene sequences by transposon insertion. In contrast, the single copy of *SIX14* present in Fola4 AJ516 and AJ705 was highly upregulated *in planta*. Loss of *SIX14* expression by pseudogenesis for all copies in Fola1 is a potential key difference between the Fola races that could contribute to their differential virulence on lettuce cultivars. Currently, there is no information on target sites or interacting proteins for *SIX14* although Yu et al (2014) placed *SIX14* in ‘effector family 4’ alongside *SIX5* and *PSE1/PSL1*, based on structural similarity of the proteins. However, unlike other members of this group, which are divergently transcribed with another paired effector, Fola *SIX14* does not appear to have a similarly paired transcribed gene. An important finding was that all Fola4 isolates contained a single copy of *SIX8* which was highly induced *in planta* but was absent in Fola1. Two different sequence variations of *SIX8* were also identified within multiple Fola4 isolates, which contributed with other evidence to the conclusion that there are two Fola4 variants. *SIX8* has been reported to be required for virulence of *F. oxysporum* f.sp. *cubense* tropical race 4 (Focub TR4) on Cavendish banana and is absent in Focub race1 (An et al., 2019) and interestingly, two sequence variants of *SIX8* have been identified that separate Focub TR4 from subtropical race 4 (Fraser-Smith et al., 2014). *SIX8* has also been shown to be required for virulence of Focn (isolate cong1-1) on *Arabidopsis* where it may also interact with the divergently transcribed effector PSE1 (paired with *SIX8*; Ayukawa et al., 2021). More recently, evidence has suggested that two specific members of the TOPLESS gene family in tomato act as susceptibility factors via interaction with SIX8 (Aalders et al., 2023).

RNAseq of Fola1 and Fola4 during lettuce infection identified both shared and race specific expressed putative effectors in addition to the *SIX* genes that require further investigation and characterisation to determine their potential role in pathogenicity. The majority of these corresponded to either candidate effector clusters (CECs) identified using the *mimp*-based effector discovery pipeline (Supplementary data SD2) or to putative effectors identified in other *F. oxysporum* f.spp. Of the 25 putative effectors identified as highly expressed *in planta* across all Fola isolates, 12 corresponded to CECs identified using the effector pipeline.

Five of these 25 putative effectors were not within 2.5kb of a *mimp*, and of these, two (OG0000220, OG0001047) corresponded to CECs; given the way the pipeline was implemented, this implied that these were orthologues of effectors in other *F. oxysporum* f.spp. that were associated with a *mimp*. The lack of any DNA or protein sequence similarity between effectors in *F. oxysporum* emphasises the importance of understanding effector protein structure (Yu et al., 2023) and highlighted some short-comings of the *mimp*-based effector discovery pipeline employed here. *Mimp*-associated candidate effectors from each *F. oxysporum* genome were clustered at 65% identity using CD-HIT which designates the longest sequence within each cluster as the representative sequence, which is then submitted to EffectorP for identification. Consequently, if the representative sequence does not meet the EffectorP probability threshold for effector identification, other sequences within the cluster which do are discarded with the entire CEC and this was the case for *SIX14*, *SIX8*, and *PSE1*. This highlights the importance of RNAseq expression data to further inform genome and effectorome analyses to enable more reliable identification of expressed putative effectors.

### Fola4 consists of two different variant forms

This study importantly presented evidence that there are two Fola4 variants represented by Fola4 isolate AJ516 and Fola4 isolates AJ705 / AJ592. Firstly, the phylogeny of both core and accessory genomes showed that Fola4 AJ516 diverges from isolates AJ705 and AJ592. Synteny analysis of the accessory contigs then demonstrated the extremely high similarity between Fola4 AJ705 and AJ592 while in contrast, although a few Fola4 AJ516 accessory genome contigs shared large blocks of synteny with AJ705 contigs, there were many rearrangements and regions of little or no synteny between the two isolates. This is consistent with the Fola4 isolates sharing a common origin with each of the variants arising separately. This conclusion is also supported by evidence from genotyping-by-sequencing data for Fola1 and Fola 4 which also suggested two introductions of Fola4 (Claerbout et al., 2022). Fola 4 isolate variation was also supported by the isolate AJ516 *SIX8-PSL1* gene pair showing significant differences compared with the *SIX8-PSE1* pair in Fola4 isolates AJ705 / AJ592. Sanger sequencing of this region showed that the *SIX8-PSE1* and *SIX8-PSL1* sequence differences were consistent across 13 Fola4 isolates (data not shown), partitioning them into two variant groups. Of particular interest is the 10 amino acid difference at the C-terminus of PSE1 (AJ705 / AJ592) compared to PSL1 (AJ516). *PSE1* has been identified in Focn isolates virulent on *Arabidopsis* while *PSL1* was found in the tomato-infecting isolate Fol 4287 (Ayukawa et al., 2021). It was then demonstrated that PSL1 could not be substituted for PSE1 in Focn mutants to confer virulence on *Arabidopsis* (Ayukawa et al., 2021). The authors therefore suggested that *PSE* / *PSL1* might play a role in host specificity, although no evidence of a direct interaction between SIX8 and PSE1 proteins was found in a yeast 2-hybrid assay. More recently however, Yu et al (2024) demonstrated a direct interaction between SIX8-PSE1 proteins *in vitro,* by co-incubation of expressed proteins and size exclusion chromatography. Given this information, the difference in the *SIX8-PSE1 / PSL1* pairs in the two Fola4 variants suggests that they could differ in host range, potentially affecting different lettuce cultivars or even different host species. This hypothesis is supported by a preliminary study in Belgium where a ‘new’ Fola4 isolate (designated FOL4+) was found to be virulent on lettuce cultivars resistant to a standard Fola4 isolate (Dockx et al., 2023). The two Fola4 variants might also have gained the *SIX8-PSE1/PSL1* gene pairs from different sources. In the Fola4 AJ705 / AJ592 variant, the presence of the *SIX8-PSE1* gene pair is close to the *SIX9.1* gene copy on a small accessory chromosome (contig20, 1.12 Mb), which may suggest that this variant has acquired these versions of *SIX8, PSE1* and *SIX9* together by HCT. In contrast, the Fola4 AJ516 variant may have gained the *SIX8-PSL1* pair by the involvement of (identical) FoHeli1-like helitrons that flank a 39 kb block containing the *SIX8-PSL1* locus. This may have been either by helitron insertion of the block following gene capture or by recombination with a helitron already present. Helitrons replicate by a rolling-circle mechanism and are capable of capturing other genes in large fragments, sometimes containing many genes, thus facilitating movement, rearrangement or duplication of genes (Thomas and Pritham, 2015; Barro-Trastoy and Kohler, 2024). Five classes of non-canonical helitrons with distinct end terminal sequences have been identified in the FOSC, with both autonomous and non-autonomous elements present in varying numbers and evidence of circular intermediates suggesting active helitron elements (Chellapan et al., 2016). Helitrons have also been shown to affect gene expression (Castanera et al., 2014) and were implicated in the loss of *SIX4* (*Avr1*) which resulted in a new race of Fol by homologous recombination (Biju et al., 2017). Transfer of chromosomes and partial chromosomes between members of the FOSC has also been demonstrated (Ma et al., 2010; Vlardingerbroek et al., 2016; van Dam et al., 2017; Li et al., 2020; Henry et al., 2021) and there is also evidence for the transfer of genes or blocks of genes (Simbaqueba et al, 2018; Liu et al., 2019). The mechanism by which this happens is not clear and needs further investigation but the involvement of recombination, or transposon (including helitrons) activity is implicated. The FoHeli1-like helitrons flanking the 39 kb block in Fola4 AJ516 that contains the *SIX8-PSL1* locus also appear to be currently active. A BLAST search of the Fola genomes revealed that other than these three identical copies, Fola4 AJ516 contains 79 additional copies of this FoHeli1-like helitron that differ by a single synonymous SNP, and a further eight copies with a non-synonymous SNP, while Fola4 AJ705 / AJ592 contains 4 identical copies, and a further 80 copies with the same synonymous SNP. In contrast, Fola1 AJ520 has only 16 identical copies of this FoHeli1-like helitron and another 3 with single SNPs, Focn 5176 contains 28 and Fol 4287 has no identical copies. Other FoHeli helitron variants are present in both Fola1 and Fola4 genomes and require a further, more detailed study to understand their involvement in genome reorganisation and the evolution of Fola races. Finally, in addition to the above differences between the two Fola4 variants, there is further provisional evidence of HCT in Fola4 AJ516 that is not present in Fola4 AJ705 / AJ592. Accessory genome contigs 21 and 28 in Fola4 AJ516 each have a telomere and, across their length, show very low levels of homology to AJ705. The subtelomere sequences of these two contigs in Fola4 AJ516 were identical and unique while all other Fola4 subtelomere sequences we were able to assemble fell into a single clade. This suggests that contigs 21 and 28 in Fola4 AJ516 might be a recent acquisition from an unknown source.

Interestingly, a preliminary search for similarity to other subtelomeric sequences from other Nanopore genome assemblies, found that these two AJ516 contig subtelomere sequences grouped very closely with those from *F. oxysporum* rocket AJ174 (data not shown). With more high quality Nanopore assemblies becoming available, the number of full-length accessory chromosomes or large contigs containing telomeric sequences will enable a more thorough analysis of these regions and potentially increase our understanding of HCT between members of the FOSC.

In summary we have presented the first comparative genome and transcriptome analyses of two globally important Fola races and presented evidence for independent evolution of Fola1 and Fola4 and for two Fola4 variant forms. In addition, we identified some key putative effectors that are highly upregulated *in planta*, which are common to both races, as well as others that are potentially important in differentiating between Fola1 and Fola4, hence providing essential information for future functional studies. We also provided evidence of potential HCT in each of the two Fola4 variants, as well as the potential involvement of helitrons in affecting gene movement and expression, all of which suggests that Fola is rapidly evolving. A comprehensive genomics study of multiple Fola isolates from all four known races would enable even greater insights into race evolution and a much better understanding of the potential role of HCT and helitrons in driving this process.

## Data Availability Statement

The datasets generated for this study can be found in the article and supplementary material. All raw sequencing data and new genome assemblies presented have been submitted to the NCBI under the BioProject ID PRJNA1092066. Further enquiries can be directed to the corresponding author.

## Author Contributions

HB: Conceptualisation, Investigation, Formal analysis, Methodology, Validation, Visualisation, Project administration, Writing – original draft, Writing - review and editing. JP: Investigation, Formal analysis, Methodology, Software, Visualisation, Writing – original draft, Writing - review and editing. RJP: Formal analysis, Visualisation, Data curation, Writing – original draft, review and editing. SJ: Investigation,Writing – original draft, Writing - review and editing JC: Formal analysis, Visualisation, Writing – original draft, Writing - review and editing AA: Formal analysis, Writing – review and edit, AL: Investigation, Writing - review and editing RJH: Conceptualisation, Funding acquisition, Supervision, Writing – original draft, Writing – review and edit, JPC: Conceptualisation, Funding acquisition, Project administration, Supervision, Writing – original draft, Writing – review and edit..

## Acknowledgments

We very gratefully acknowledge Giovanna Gilardi (University of Turin), Mathieu Pel (Enza Zaden, NL), Bart Geraats (BASF, NL) and G’s Espana for supplying isolates. We also thank Laura Baxter (University of Warwick) for advice on RNAseq analysis.

## References

Aalders, T.R., de Sain, M., Gawehns, F., Oudejans, N., Jak, Y.D., Dekker, H.L., Rep, M., van den Burg, H.A., and Takken, F.L.W. (2024). Specific members of the TOPLESS family are susceptibility genes for Fusarium wilt in tomato and Arabidopsis. Plant Biotechnology Journal 22(1), 248–261. doi: 10.1111/pbi.14183.

Aimé, S., Alabouvette, C., Steinberg, C., and Olivain, C. (2013). The endophytic strain *Fusarium oxysporum* Fo47: a good candidate for priming the defense responses in tomato roots. Molecular Plant-Microbe Interactions 26(8), 918–926. doi: 10.1094/MPMI-12-12-0290-R

Almagro Armenteros, J.J., Tsirigos, K.D., Sønderby, C.K., Petersen, T.N., Winther, O., Brunak, S., von Heijne, G., and Nielsen, H. (2019). SignalP 5.0 improves signal peptide predictions using deep neural networks. Nature Biotechnology 37(4), 420–423. doi: 10.1038/s41587-019-0036-z

An, B., Hou, X., Guo, Y., Zhao, S., Luo, H., He, C., and Wang, Q. (2019). The effector SIX8 is required for virulence of *Fusarium oxysporum* f.sp. *cubense* tropical race 4 to Cavendish banana. Fungal Biology 123(5), 423–430. doi: 10.1016/j.funbio.2019.03.001.

Armitage, A.D., Cockerton, H.M., Sreenivasaprasad, S., Woodhall, J., Lane, C.R., Harrison, R.J., and Clarkson, J.P. (2020). Genomics, evolutionary history and diagnostics of the *Alternaria alternata* species group including apple and asian pear pathotypes. Frontiers in Microbiology 10. doi: 10.3389/fmicb.2019.03124.

Armitage, A.D., Taylor, A., Sobczyk, M.K., Baxter, L., Greenfield, B.P.J., Bates, H.J., Wilson, F., Jackson, A.C., Ott, S., and Harrison, R.J. (2018). Characterisation of pathogen-specific regions and novel effector candidates in *Fusarium oxysporum* f.sp. *cepae*. Scientific Reports 8(1), 13530. doi: 10.1038/s41598-018-30335-7.

Aronesty, E. (2013). Comparison of sequencing utility programs. The Open Bioinformatics Journal 7(1). Doi: 10.2174/1875036201307010001

Asai, S., Ayukawa, Y., Gan, P., and Shirasu, K. (2021). Draft Genome Resources for Brassicaceae Pathogens *Fusarium oxysporum* f. sp. *raphani* and *Fusarium oxysporum* f.sp. *rapae*. Molecular Plant-Microbe Interactions 34(11), 1316–1319. Doi: 10.1094/MPMI-06-21-0148-A

Ayukawa, Y., Asai, S., Gan, P., Tsushima, A., Ichihashi, Y., Shibata, A., Komatsu, K., Houterman, P.M., Rep, M., and Shirasu, K. (2021). A pair of effectors encoded on a conditionally dispensable chromosome of *Fusarium oxysporum* suppress host-specific immunity. Communications Biology 4(1), 707. Doi: 10.1038/s42003-021-02245-4

Barro-Trastoy, D., and Köhler, C. (2024). Helitrons: genomic parasites that generate developmental novelties. Trends in Genetics 0(0). doi: 10.1016/j.tig.2024.02.002.

Bendtsen, J.D., Nielsen, H., Von Heijne, G., and Brunak, S. (2004). Improved prediction of signal peptides: SignalP 3.0. Journal of Molecular Biology 340(4), 783–795. doi: 10.1016/j.jmb.2004.05.028

Biju, V.C., Fokkens, L., Houterman, P.M., Rep, M., and Cornelissen, B.J.C. (2017). Multiple evolutionary trajectories have led to the emergence of races in *Fusarium oxysporum* f.sp. *lycopersici*. Applied and Environmental Microbiology 83(4), e02548–02516. doi: 10.1128/AEM.02548-16.

Blin, K., Shaw, S., Kloosterman, A.M., Charlop-Powers, Z., Van Wezel, G.P., Medema, M.H., and Weber, T. (2021). antiSMASH 6.0: improving cluster detection and comparison capabilities. Nucleic Acids Research 49(W1), W29–W35. doi: 10.1093/nar/gkab335

Bray, N.L., Pimentel, H., Melsted, P., and Pachter, L. (2016). Near-optimal probabilistic RNA-seq quantification. Nature Biotechnology 34(5), 525–527. doi: 10.1038/nbt.3519

Brenes Guallar, M.A., Fokkens, L., Rep, M., Berke, L., and van Dam, P. (2022). *Fusarium oxysporum* effector clustering version 2: An updated pipeline to infer host range. Frontiers in Plant Science 13, 1012688. doi: 10.3389/fpls.2022.1012688

Brůna, T., Hoff, K.J., Lomsadze, A., Stanke, M., and Borodovsky, M. (2021). BRAKER2: automatic eukaryotic genome annotation with GeneMark-EP+ and AUGUSTUS supported by a protein database. NAR Genomics and Bioinformatics 3(1), 108. doi: 10.1093/nargab/iqaa108

Camacho, C., Coulouris, G., Avagyan, V., Ma, N., Papadopoulos, J., Bealer, K., and Madden, T.L. (2009). BLAST+: architecture and applications. BMC ioinformatics 10, 1–9. Doi: 10.1186/1471-2105-10-421

Castanera, R., Pérez, G., López, L., Sancho, R., Santoyo, F., Alfaro, M., Gabaldón, T., Pisabarro, A.G., Oguiza, J.A., and Ramírez, L. (2014). Highly expressed captured genes and cross-kingdom domains present in helitrons create novel diversity in *Pleurotus ostreatus* and other fungi. BMC Genomics 15(1), 1071. doi: 10.1186/1471-2164-15-1071.

Chang, W., Li, H., Chen, H., Qiao, F., and Zeng, H. (2020). Identification of *mimp*-associated effector genes in *Fusarium oxysporum* f. sp. *cubense* race 1 and race 4 and virulence confirmation of a candidate effector gene. Microbiological Research 232, 126375. doi: 10.1016/j.micres.2019.126375.

Chellapan, B.V., van Dam, P., Rep, M., Cornelissen, B.J.C., and Fokkens, L. (2016). Non-canonical helitrons in *Fusarium oxysporum*. Mobile DNA 7(1), 27. doi: 10.1186/s13100-016-0083-7.

Chen, Y., Nie, F., Xie, S.-Q., Zheng, Y.-F., Dai, Q., Bray, T., Wang, Y.-X., Xing, J.-F., Huang, Z.-J., Wang, D.-P., He, L.-J., Luo, F., Wang, J.-X., Liu, Y.-Z., and Xiao, C.-L. (2021). Efficient assembly of nanopore reads via highly accurate and intact error correction. Nature Communications 12(1), 60. doi: 10.1038/s41467-020-20236-7.

Chiara, M., Fanelli, F., Mulè, G., Logrieco, A.F., Pesole, G., Leslie, J.F., Horner, D.S., and Toomajian, C. (2015). Genome sequencing of multiple isolates highlights subtelomeric genomic diversity within *Fusarium fujikuroi*. Genome Biology and Evolution 7(11), 3062–3069. doi: 10.1093/gbe/evv198.

Claerbout, J., Van Poucke, K., Mestdagh, H., Delaere, I., Vandevelde, I., Venneman, S., Decombel, A., Bleyaert, P., Neukermans, J., Viaene, N., Heungens, K., and Höfte, M. (2023). *Fusarium* isolates from Belgium causing wilt in lettuce show genetic and pathogenic diversity. Plant Pathology 72(3), 593–609. doi: 10.1111/ppa.13668.

Claerbout, J., Venneman, S., Vandevelde, I., Decombel, A., Bleyaert, P., Volckaert, A., Neukermans, J., and Höfte, M. (2018). First report of *Fusarium oxysporum* f.sp. lactucae race 4 on lettuce in Belgium. Plant Disease 102(5), 1037. doi: 10.1094/pdis-10-17-1627-pdn.

Constantin, M.E., Fokkens, L., de Sain, M., Takken, F.L.W., and Rep, M. (2021). Number of candidate effector genes in accessory genomes differentiates pathogenic from endophytic *Fusarium oxysporum* strains. Frontiers in Plant Science 12, 761740. doi: 10.3389/fpls.2021.761740

Czislowski, E., Fraser-Smith, S., Zander, M., O’Neill, W.T., Meldrum, R.A., Tran-Nguyen, L.T.T., Batley, J., and Aitken, E.A.B. (2018). Investigation of the diversity of effector genes in the banana pathogen, Fusarium oxysporum f.sp. cubense, reveals evidence of horizontal gene transfer. Molecular Plant Pathology 19(5), 1155–1171. doi: 10.1111/mpp.12594

De Coster, W., D’Hert, S., Schultz, D.T., Cruts, M., and Van Broeckhoven, C. (2018). NanoPack: visualizing and processing long-read sequencing data. Bioinformatics 34(15), 2666–2669. doi: 10.1093/bioinformatics/bty149.

Dobin, A., Davis, C.A., Schlesinger, F., Drenkow, J., Zaleski, C., Jha, S., Batut, P., Chaisson, M., and Gingeras, T.R. (2013). STAR: ultrafast universal RNA-seq aligner. Bioinformatics 29(1), 15–21. doi: 10.1093/bioinformatics/bts635

Dockx, T., Mestdagh, H., Arnouts, T., Van Mullem, J., Vandevelde, I., Höfte, M. and Heungens, K. (2023). ‘Vooral lollo bionda, lollo rossa en Romeinse sla tolerant op grond met Fol 4+’. Proeftuinnieuws 15, 12–13.

Edel-Hermann, V., and Lecomte, C. (2019). Current status of *Fusarium oxysporum formae speciales* and races. Phytopathology 109(4), 512–530. doi: 10.1094/phyto-08-18-0320-rvw.

Emms, D.M., and Kelly, S. (2019). OrthoFinder: phylogenetic orthology inference for comparative genomics. Genome biology 20, 1–14. doi: 10.1186/s13059-019-1832-y

Finn, R.D., Clements, J., and Eddy, S.R. (2011). HMMER web server: interactive sequence similarity searching. Nucleic Acids Research 39, W29–W37. doi: 10.1093/nar/gkr367

Fokkens, L., Guo, L., Dora, S., Wang, B., Ye, K., Sánchez-Rodríguez, C., and Croll, D. (2020). A chromosome-scale genome assembly for the *Fusarium oxysporum* strain Fo5176 to establish a model *Arabidopsis*-fungal pathosystem. G3: Genes, Genomes, Genetics 10(10), 3549–3555.

Fraser-Smith, S., Czislowski, E., Meldrum, R.A., Zander, M., O’Neill, W., Balali, G.R., and Aitken, E.A.B. (2014). Sequence variation in the putative effector gene SIX8 facilitates molecular differentiation of *Fusarium oxysporum* f. sp. *cubense*. Plant Pathology 63(5), 1044–1052. doi: 10.1111/ppa.12184.

Fu, L., Niu, B., Zhu, Z., Wu, S., and Li, W. (2012). CD-HIT: accelerated for clustering the next-generation sequencing data. Bioinformatics 28(23), 3150–3152. doi: 10.1093/bioinformatics/bts565

Gálvez, L., Brizuela, A.M., Garcés, I., Cainarca, J.S., and Palmero, D. (2023). First report of *Fusarium oxysporum* f.sp. *lactucae* race 4 causing lettuce wilt in Spain. Plant Disease 107 (8), 2549. doi: 10.1094/pdis-12-22-2819-pdn.

Gao, W., Khang, C.H., Park, S.-Y., Lee, Y.-H., and Kang, S. (2002). Evolution and organization of a highly dynamic, subtelomeric helicase gene family in the rice blast fungus *Magnaporthe grisea*. Genetics 162(1), 103–112. doi: 10.1093/genetics/162.1.103.

Garibaldi, A., Gilardi, G., and Gullino, M.L. (2004). Varietal resistance of lettuce to *Fusarium oxysporum* f.sp. *lactucae*. Crop Protection 23(9), 845–851. doi: 10.1016/j.cropro.2004.01.005.

Gilardi, G., Franco Ortega, S., Van Rijswick, P.C.J., Ortu, G., Gullino, M.L., and Garibaldi, A. (2017a). A new race of *Fusarium oxysporum* f.sp. *lactucae* of lettuce. Plant Pathology 66(4), 677–688. doi: 10.1111/ppa.12616

Gilardi, G., Garibaldi, A., Matic, S., Senatore, M.T., Pipponzi, S., Prodi, A., and Gullino, M.L. (2019). First report of *Fusarium oxysporum* f.sp. *lactucae* race 4 on lettuce in Italy. Plant Disease 103(10), 2680–2680. doi: 10.1094/pdis-05-19-0902-pdn.

Gilardi, G., Pons, C., Gard, B., Franco-Ortega, S., and Gullino, M.L. (2017b). Presence of Fusarium wilt, incited by *Fusarium oxysporum* f.sp. *lactucae*, on lettuce in France. Plant Disease 101(6), 1053. doi: 10.1094/PDIS-12-16-1815-PDN

Gilardi, G., Vasileiadou, A., Garibaldi, A., and Gullino, M.L. (2021). Low temperatures favour Fusarium wilt development by race 4 of *Fusarium oxysporum* f.sp. *lactucae*. Journal of Plant Pathology 103(3), 973–979. doi: 10.1007/s42161-021-00859-5.

Grabherr, M.G., Russell, P., Meyer, M., Mauceli, E., Alföldi, J., Di Palma, F., and Lindblad-Toh, K. (2010). Genome-wide synteny through highly sensitive sequence alignment: Satsuma. Bioinformatics 26(9), 1145–1151. doi: 10.1093/bioinformatics/btq102

Guerrero, M.M., Martínez, M.C., León, M., Armengol, J., and Monserrat, A. (2020). First report of Fusarium wilt of lettuce caused by *Fusarium oxysporum* f.sp. *lactucae* race 1 in Spain. Plant Disease 104(6), 1858–1858. doi: 10.1094/pdis-10-19-2143-pdn.

Guo, L., Han, L., Yang, L., Zeng, H., Fan, D., Zhu, Y., Feng, Y., Wang, G., Peng, C., and Jiang, X. (2014). Genome and transcriptome analysis of the fungal pathogen *Fusarium oxysporum* f. sp. *cubense* causing banana vascular wilt disease. PLoS One 9(4), e95543. doi: 10.1371/journal.pone.0095543

Gupta, S., Gallavotti, A., Stryker, G.A., Schmidt, R.J., and Lal, S.K. (2005). A novel class of helitron-related transposable elements in maize contain portions of multiple pseudogenes. Plant Molecular Biology 57(1), 115–127. doi: 10.1007/s11103-004-6636-z.

He, W., Yang, J., Jing, Y., Xu, L., Yu, K., and Fang, X. (2023). NGenomeSyn: an easy-to-use and flexible tool for publication-ready visualization of syntenic relationships across multiple genomes. Bioinformatics 39(3), btad121. doi: 10.1093/bioinformatics/btad121.

Henry, P., Kaur, S., Pham, Q.A.T., Barakat, R., Brinker, S., Haensel, H., Daugovish, O., and Epstein, L. (2020). Genomic differences between the new *Fusarium oxysporum* f.sp. *apii* (Foa) race 4 on celery, the less virulent Foa races 2 and 3, and the avirulent on celery f. sp. *coriandrii*. BMC Genomics 21(1), 1–23. doi: 10.1186/s12864-020-07141-5

Henry, P.M., Pincot, D.D.A., Jenner, B.N., Borrero, C., Aviles, M., Nam, M.-H., Epstein, L., Knapp, S.J., and Gordon, T.R. (2021). Horizontal chromosome transfer and independent evolution drive diversification in *Fusarium oxysporum* f.sp. *fragariae*. The New Phytologist 230(1), 327–340. doi: 10.1111/nph.17141.

Herrero, M.L., Nagy, N.E., and Solheim, H. (2021). First Report of *Fusarium oxysporum* f.sp. *lactucae* race 1 causing Fusarium wilt of lettuce in Norway. Plant Disease 105(8), 2239. doi: 10.1094/pdis-01-21-0134-pdn.

Houterman, P.M., Speijer, D., Dekker, H.L., de Koster, C.G., Cornelissen, B.J.C., and Rep, M. (2007). The mixed xylem sap proteome of *Fusarium oxysporum*-infected tomato plants. Molecular Plant Pathology 8(2), 215–221. doi: 10.1111/j.1364-3703.2007.00384.x.

Huang, X. (2023). "Host-specific subtelomere: structural variation and horizontal transfer in asexual filamentous fungal pathogens". Preprint bioRxiv. doi: 10.1101/2023.02.05.527183

Hudson, O., Fulton, J.C., Dong, A.K., Dufault, N.S., and Ali, M.E. (2021). *Fusarium oxysporum* f. sp. *niveum* molecular diagnostics past, present and future. International Journal of Molecular Sciences 22(18), 9735. doi: 10.3390/ijms22189735

Inglis, P.W., Rigden, D.J., Mello, L.V., Louis, E.J., and Valadares-Inglis, M.C. (2005). Monomorphic subtelomeric DNA in the filamentous fungus, Metarhizium anisopliae,contains a RecQ helicase-like gene. Molecular genetics and genomics: MGG 274(1), 79–90. doi: 10.1007/s00438-005-1154-5.

Jangir, P., Mehra, N., Sharma, K., Singh, N., Rani, M., and Kapoor, R. (2021). Secreted in xylem genes: Drivers of host adaptation in *Fusarium oxysporum*. Frontiers in Plant Science 12, 628611. doi: 10.3389/fpls.2021.628611

Jobe, T.O., Urner, M., Ulloa, M., Broders, K., Hutmacher, R.B., and Ellis, M.L. (2024). *Secreted in xylem* (*SIX*) gene *SIX9* is highly conserved in *Fusarium oxysporum* f. sp. *vasinfectum* race 4 isolates from cotton in the United States. PhytoFrontiers. doi: 10.1094/PHYTOFR-11-23-0143-SC.

Jones, P., Binns, D., Chang, H.-Y., Fraser, M., Li, W., McAnulla, C., McWilliam, H., Maslen, J., Mitchell, A., and Nuka, G. (2014). InterProScan 5: genome-scale protein function classification. Bioinformatics 30(9), 1236–1240. doi: 10.1093/bioinformatics/btu031

Kapitonov, V.V., and Jurka, J. (2007). Helitrons on a roll: eukaryotic rolling-circle transposons. Trends in Genetics: TIG 23(10), 521–529. doi: 10.1016/j.tig.2007.08.004.

Katoh, K., Rozewicki, J., and Yamada, K.D. (2019). MAFFT online service: multiple sequence alignment, interactive sequence choice and visualization. Briefings in Bioinformatics 20(4), 1160–1166. doi: 10.1093/bib/bbx108

Keller, O., Kollmar, M., Stanke, M., and Waack, S. (2011). A novel hybrid gene prediction method employing protein multiple sequence alignments. Bioinformatics 27(6), 757–763. doi: 10.1093/bioinformatics/btr010

Kolde, R., and Kolde, M.R. (2015). Package ‘pheatmap’. R package 1(7), 790.

Krasnov, G.S., Pushkova, E.N., Novakovskiy, R.O., Kudryavtseva, L.P., Rozhmina, T.A., Dvorianinova, E.M., Povkhova, L.V., Kudryavtseva, A.V., Dmitriev, A.A., and Melnikova, N.V. (2020). High-quality genome assembly of *Fusarium oxysporum* f. sp. *lini*. Frontiers in genetics 11, 959. doi: 10.3389/fgene.2020.00959

Krzywinski, M., Schein, J., Birol, I., Connors, J., Gascoyne, R., Horsman, D., Jones, S.J., and Marra, M.A. (2009). Circos: an information aesthetic for comparative genomics. Genome Research 19(9), 1639–1645. doi: 10.1101/gr.092759.109

Langmead, B., and Salzberg, S.L. (2012). Fast gapped-read alignment with Bowtie 2. Nature Methods 9(4), 357–359. doi: 10.1038/nmeth.1923

Letunic, I., and Bork, P. (2021). Interactive Tree Of Life (iTOL) v5: an online tool for phylogenetic tree display and annotation. Nucleic Acids Research 49(W1), W293–W296. doi: 10.1093/nar/gkab301

Li, H. (2013). Aligning sequence reads, clone sequences and assembly contigs with BWA-MEM. arXiv preprint arXiv:1303.3997.

Li, H., Handsaker, B., Wysoker, A., Fennell, T., Ruan, J., Homer, N., Marth, G., Abecasis, G., Durbin, R., and Genome Project Data Processing, S. (2009). The sequence alignment/map format and SAMtools. Bioinformatics 25(16), 2078–2079. doi: 10.1093/bioinformatics/btp352

Li, J., Fokkens, L., Conneely, L.J., and Rep, M. (2020). Partial pathogenicity chromosomes in *Fusarium oxysporum* are sufficient to cause disease and can be horizontally transferred. Environmental Microbiology 22(12), 4985–5004. doi: 10.1111/1462-2920.15095

Li, J., Fokkens, L., van Dam, P., and Rep, M. (2020). Related mobile pathogenicity chromosomes in *Fusarium oxysporum* determine host range on cucurbits. Molecular Plant Pathology 21(6), 761–776. doi: 10.1111/mpp.12927

Liu, S., Wu, B., Lv, S., Shen, Z., Li, R., Yi, G., Li, C., and Guo, X. (2019). Genetic diversity in FUB Genes of *Fusarium oxysporum* f. sp. *cubense* suggests horizontal gene transfer. Frontiers in Plant Science 10. doi: 10.3389/fpls.2019.01069.

Love, M.I., Huber, W., and Anders, S. (2014). Moderated estimation of fold change and dispersion for RNA-seq data with DESeq2. Genome Biology 15(12), 1–21. 10.1186/s13059-014-0550-8

Ma, L.-J., Van Der Does, H.C., Borkovich, K.A., Coleman, J.J., Daboussi, M.-J., Di Pietro, A., Dufresne, M., Freitag, M., Grabherr, M., and Henrissat, B. (2010). Comparative genomics reveals mobile pathogenicity chromosomes in *Fusarium*. Nature 464(7287), 367–373. doi: 10.1038/nature08850.

McDonald, M.C., Taranto, A.P., Hill, E., Schwessinger, B., Liu, Z., Simpfendorfer, S., Milgate, A., and Solomon, P.S. (2019). Transposon-mediated horizontal transfer of the host-specific virulence protein ToxA between three fungal wheat pathogens. mBio 10(5), 10.1128/mbio.01515-01519. doi: 10.1128/mbio.01515-19.

Motohashi, S. (1960). Occurrence of lettuce root rot. Annals of the Phytopathological Society of Japan 25, 47.

Murray, J.J., Hisamutdinova, G., Sandoya, G.V., Raid, R.N., and Slinski, S. (2021). Genetic resistance of *Lactuca* spp. against *Fusarium oxysporum* f. sp. *lactucae* race 1. HortScience 56(12), 1552–1564. doi: 10.21273/hortsci16186-21.

Nguyen, L.-T., Schmidt, H.A., Von Haeseler, A., and Minh, B.Q. (2015). IQ-TREE: a fast and effective stochastic algorithm for estimating maximum-likelihood phylogenies. Molecular Biology and Evolution 32(1), 268–274. doi: 10.1093/molbev/msu300

Nielsen, H. (2017). Predicting secretory proteins with SignalP. Protein Function Prediction: Methods and Protocols, 59–73. doi: 10.1007/978-1-4939-7015-5_6

Nielsen, H., and Krogh, A. (1998). Prediction of signal peptides and signal anchors by a hidden Markov model, in: *Ismb*, 122–130.

Pasquali, M., Dematheis, F., Gullino, M.L., and Garibaldi, A. (2007). Identification of race 1 of *Fusarium oxysporum* f. sp. *lactucae* on lettuce by inter-retrotransposon sequence-characterized amplified region technique. Phytopathology 97(8), 987–996. doi: 10.1094/phyto-97-8-0987.

Petersen, T.N., Brunak, S., Von Heijne, G., and Nielsen, H. (2011). SignalP 4.0: discriminating signal peptides from transmembrane regions. Nature Methods 8(10), 785–786. doi: 10.1038/nmeth.1701

Quinlan, A.R., and Hall, I.M. (2010). BEDTools: a flexible suite of utilities for comparing genomic features. Bioinformatics 26(6), 841–842. doi: 10.1093/bioinformatics/btq033

Rahnama, M., Wang, B., Dostart, J., Novikova, O., Yackzan, D., Yackzan, A., Bruss, H., Baker, M., Jacob, H., Zhang, X., Lamb, A., Stewart, A., Heist, M., Hoover, J., Calie, P., Chen, L., Liu, J., and Farman, M.L. (2021). Telomere roles in fungal genome evolution and adaptation. Frontiers in Genetics 12. doi: 10.3389/fgene.2021.676751.

Rempel, A., and Wittler, R. (2021). SANS serif: alignment-free, whole-genome-based phylogenetic reconstruction. Bioinformatics 37(24), 4868–4870. doi: 10.1093/bioinformatics/btab444.

Salimi, S., Abdi, M.F., and Rahnama, M. (2024). Characterization and organization of telomeric-linked helicase (TLH) gene families in *Fusarium oxysporum*. Preprint BioRxiv. doi: 10.1101/2024.02.27.582403

Sánchez-Alonso, P., and Guzmán, P. (1998). Organization of chromosome ends in *Ustilago maydis* RecQ-like helicase motifs at telomeric regions. Genetics 148(3), 1043–1054. doi: 10.1093/genetics/148.3.1043.

Schmidt, S.M., Houterman, P.M., Schreiver, I., Ma, L., Amyotte, S., Chellappan, B., Boeren, S., Takken, F.L.W., and Rep, M. (2013). MITEs in the promoters of effector genes allow prediction of novel virulence genes in *Fusarium oxysporum*. BMC Genomics 14(1), 119. doi: 10.1186/1471-2164-14-119.

Scott, J.C., Kirkpatrick, S.C., and Gordon, T.R. (2010). Variation in susceptibility of lettuce cultivars to fusarium wilt caused by *Fusarium oxysporum* f.sp. *lactucae*. Plant Pathology 59(1), 139–146. doi: 10.1111/j.1365-3059.2009.02179.x.

Seo, S., Pokhrel, A., and Coleman, J.J. (2020). The genome sequence of five genotypes of *Fusarium oxysporum* f. sp. *vasinfectum*: A resource for studies on Fusarium wilt of cotton. Molecular Plant-Microbe Interactions 33(2), 138–140. doi: 10.1094/MPMI-07-19-0197-A

Simão, F.A., Waterhouse, R.M., Ioannidis, P., Kriventseva, E.V., and Zdobnov, E.M. (2015). BUSCO: assessing genome assembly and annotation completeness with single-copy orthologs. Bioinformatics 31(19), 3210–3212. doi: 10.1093/bioinformatics/btv351.

Simbaqueba, J., Catanzariti, A.M., González, C., and Jones, D.A. (2018). Evidence for horizontal gene transfer and separation of effector recognition from effector function revealed by analysis of effector genes shared between cape gooseberry- and tomato-infecting *formae speciales* of *Fusarium oxysporum*. Molecular Plant Pathology 19(10), 2302–2318. doi: 10.1111/mpp.12700.

Simbaqueba, J., Rodríguez, E.A., Burbano-David, D., González, C., and Caro-Quintero, A. (2021). Putative novel effector genes revealed by the genomic analysis of the phytopathogenic fungus *Fusarium oxysporum* f. sp. *physali* (Foph) that infects cape gooseberry plants. Frontiers in Microbiology 11. doi: 10.3389/fmicb.2020.593915.

Sperschneider, J., Gardiner, D.M., Dodds, P.N., Tini, F., Covarelli, L., Singh, K.B., Manners, J.M., and Taylor, J.M. (2016). EffectorP: predicting fungal effector proteins from secretomes using machine learning. New Phytologist 210(2), 743–761. doi: 10.1111/nph.13794

Stanke, M., Keller, O., Gunduz, I., Hayes, A., Waack, S., and Morgenstern, B. (2006). AUGUSTUS: ab initio prediction of alternative transcripts. Nucleic Acids Research 34(suppl_2), W435–W439. doi: 10.1093/nar/gkl200

Takken, F., and Rep, M. (2010). The arms race between tomato and *Fusarium oxysporum*. Molecular Plant Pathology 11(2), 309–314. doi: 10.1111/j.1364-3703.2009.00605.x

Taylor, A., Armitage, A.D., Handy, C., Jackson, A.C., Hulin, M.T., Harrison, R.J., and Clarkson, J.P. (2019a). Basal Rot of Narcissus: Understanding Pathogenicity in *Fusarium oxysporum* f. sp. *narcissi*. Frontiers in Microbiology 10. doi: 10.3389/fmicb.2019.02905.

Taylor, A., Barnes, A., Jackson, A.C., and Clarkson, J.P. (2019b). First Report of *Fusarium oxysporum* and *Fusarium redolens* causing wilting and yellowing of wild rocket (*Diplotaxis tenuifolia*) in the United Kingdom. Plant Disease 103(6), 1428–1428. doi: 10.1094/pdis-12-18-2143-pdn.

Taylor, A., Jackson, A.C., and Clarkson, J.P. (2019c). First report of *Fusarium oxysporum* f. sp. *lactucae* race 4 causing lettuce wilt in England and Ireland. Plant Disease 103(5), 1033–1033. doi: 10.1094/pdis-10-18-1858-pdn.

Taylor, A., Vágány, V., Jackson, A.C., Harrison, R.J., Rainoni, A., and Clarkson, J.P. (2016). Identification of pathogenicity-related genes in *Fusarium oxysporum* f. sp. *cepae*. Molecular Plant Pathology 17(7), 1032–1047. doi: 10.1111/mpp.12346

Testa, A.C., Hane, J.K., Ellwood, S.R., and Oliver, R.P. (2015). CodingQuarry: highly accurate hidden Markov model gene prediction in fungal genomes using RNA-seq transcripts. BMC Genomics 16, 1–12. doi: 10.1186/s12864-015-1344-4

Thomas, J., and Pritham, E.J. (2015). Helitrons, the Eukaryotic rolling-circle transposable elements. Microbiology Spectrum 3(4), doi: 10.1128/microbiolspec.mdna3-0049-2014.

Trapnell, C., Williams, B.A., Pertea, G., Mortazavi, A., Kwan, G., van Baren, M.J., Salzberg, S.L., Wold, B.J., and Pachter, L. (2010). Transcript assembly and abundance estimation from RNA-Seq reveals thousands of new transcripts and switching among isoforms. Nature Biotechnology 28(5), 511. doi: 10.1038/nbt.1621

van Amsterdam, S., Jenkins, S., and Clarkson, J.P. (2023). First report of *Fusarium oxysporum* f. sp. *lactucae* race 1 causing lettuce wilt in Northern Ireland. Plant Disease 107(8), 2524.. doi: 10.1094/pdis-01-23-0196-pdn.

van Dam, P., de Sain, M., Ter Horst, A., van der Gragt, M., and Rep, M. (2018). Use of comparative genomics-based markers for discrimination of host specificity in *Fusarium oxysporum*. Applied and Environmental Microbiology 84(1), e01868–01817. doi: 10.1128/AEM.01868-17

van Dam, P., Fokkens, L., Ayukawa, Y., van der Gragt, M., Ter Horst, A., Brankovics, B., Houterman, P.M., Arie, T., and Rep, M. (2017). A mobile pathogenicity chromosome in *Fusarium oxysporum* for infection of multiple cucurbit species. Scientific Reports 7(1), 9042. doi: 10.1038/s41598-017-07995-y

van Dam, P., Fokkens, L., Schmidt, S.M., Linmans, J.H.J., Kistler, H.C., Ma, L.-J., and Rep, M. (2016). Effector profiles distinguish formae speciales of *Fusarium oxysporum*. Environmental Microbiology 18(11), 4087–4102. doi: 10.1111/1462-2920.13445

van Dam, P., and Rep, M. (2017). The distribution of miniature impala elements and *SIX* genes in the *Fusarium* genus is suggestive of horizontal gene transfer. Journal of Molecular Evolution 85(1), 14–25. doi: 10.1007/s00239-017-9801-0.

Vaser, R., Sović, I., Nagarajan, N., Šikić, M. (2017) Fast and accurate de novo genome assembly from long uncorrected reads. Genome Research 27(5), 737–746. doi: 10.1101/gr.214270.116.

Vlaardingerbroek, I., Beerens, B., Rose, L., Fokkens, L., Cornelissen, B.J.C., and Rep, M. (2016). Exchange of core chromosomes and horizontal transfer of lineage-specific chromosomes in *Fusarium oxysporum*. Environmental Microbiology 18(11), 3702–3713. doi: 10.1111/1462-2920.13281.

Vlaardingerbroek, I., Beerens, B., Schmidt, S.M., Cornelissen, B.J.C., and Rep, M. (2016). Dispensable chromosomes in *Fusarium oxysporum* f. sp. *lycopersici*. Molecular Plant Pathology 17(9), 1455–1466. doi: 10.1111/mpp.12440

Walker, B.J., Abeel, T., Shea, T., Priest, M., Abouelliel, A., Sakthikumar, S., Cuomo, C.A., Zeng, Q., Wortman, J., and Young, S.K. (2014). Pilon: an integrated tool for comprehensive microbial variant detection and genome assembly improvement. PLoS One 9(11), e112963. doi: 10.1371/journal.pone.0112963

Wang, B., Yu, H., Jia, Y., Dong, Q., Steinberg, C., Alabouvette, C., Edel-Hermann, V., Kistler, H.C., Ye, K., and Ma, L.-J. (2020). Chromosome-scale genome assembly of *Fusarium oxysporum* strain Fo47, a fungal endophyte and biocontrol agent. Molecular Plant-Microbe Interactions 33(9), 1108–1111. doi: 10.1094/MPMI-05-20-0116-A

Wang, Y., Zhao, Y., Bollas, A., Wang, Y., and Au, K.F. (2021). Nanopore sequencing technology, bioinformatics and applications. Nature Biotechnology 39(11), 1348–1365. doi: 10.1038/s41587-021-01108-x

Wheeler, T.J., and Eddy, S.R. (2013). nhmmer: DNA homology search with profile HMMs. Bioinformatics 29(19), 2487–2489. doi: 10.1093/bioinformatics/btt403

Xingxing, P., Khan, R.A.A., Yan, L., Yuhong, Y., Bingyan, X., Zhenchuan, M., and Jian, L. (2021). Draft genome resource of *Fusarium oxysporu*m f. sp. *capsici*, the infectious agent of pepper Fusarium wilt. Molecular Plant-Microbe Interactions 34(6), 715–717. doi: 10.1094/MPMI-12-20-0355-A

Yang, H., Yu, H., and Ma, L.-J. (2020). Accessory chromosomes in *Fusarium oxysporum*. Phytopathology 110(9), 1488–1496. doi: 10.1094/phyto-03-20-0069-ia.

Zhang, H., Yohe, T., Huang, L., Entwistle, S., Wu, P., Yang, Z., Busk, P.K., Xu, Y., and Yin, Y. (2018). dbCAN2: a meta server for automated carbohydrate-active enzyme annotation. Nucleic Acids Research 46(W1), W95–W101. doi: 10.1093/nar/gky418

