## Supplementary Figures for "Comparative genomics and transcriptomics reveal differences in effector complement and expression between races of *Fusarium oxysporum* f.sp. *lactucae*"

### 1 Supplementary Figure S1

**Figure S1**. Collinearity analysis of core genome sequences between *Fusarium oxysporum* f.sp *lycopersici* (Fol) isolate 4287 and *F. oxysporum* f.sp. *lactucae* (Fola) race 1 and 4 isolates. Syntenic alignments larger than 100 Kb to each Fol 4287 chromosome are highlighted in the same colour. Chr numbers represent Fol 4287 chromosomes and c numbers represent Fola contigs.

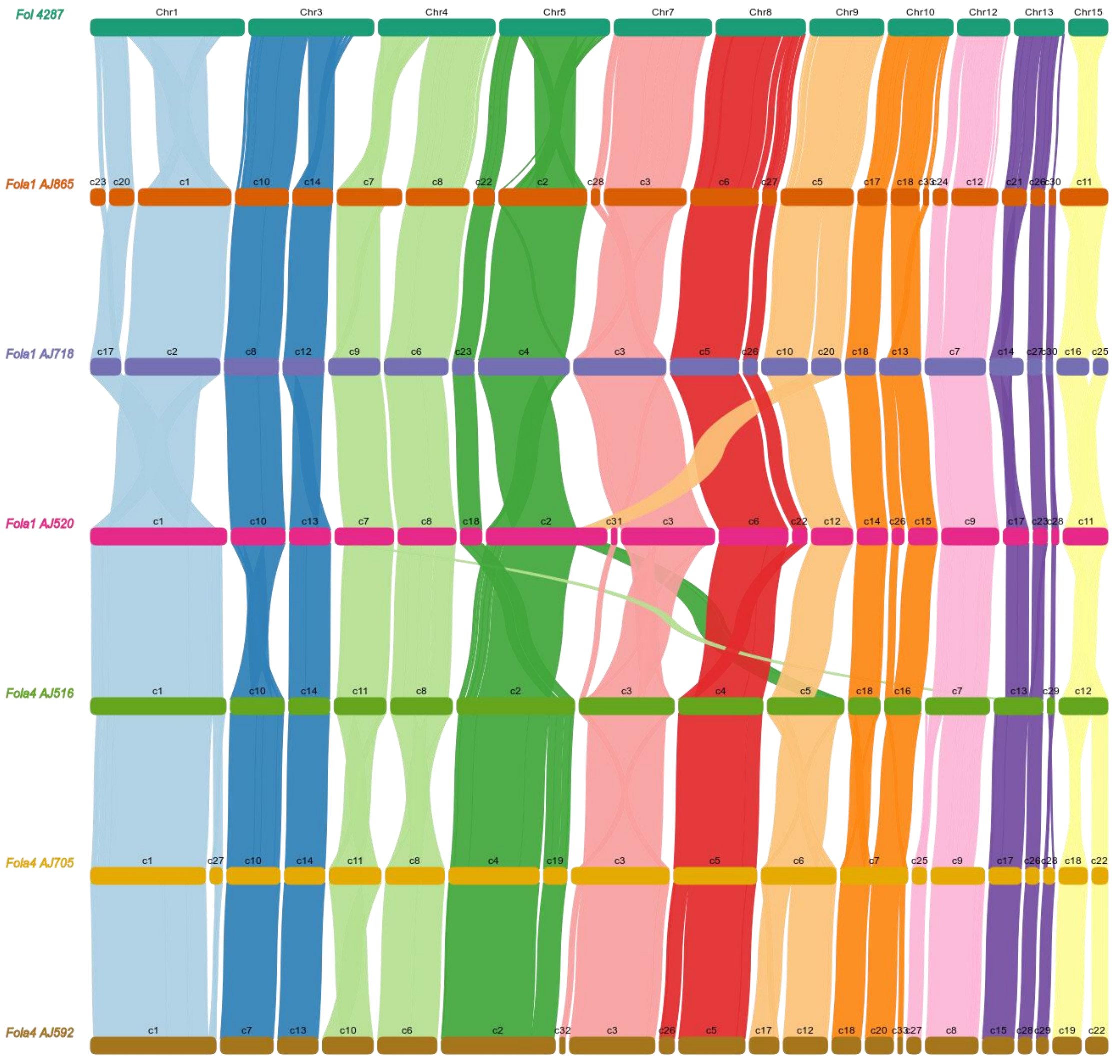

### 2 Supplementary Figure S2

**Figure S2**. Phylogeny of sub-telomeric 7.5kb sequences for *Fusarium oxysporum* f.sp. *lactucae* (Fola) race 1 and 4 isolates. Fola4 AJ516 contigs 21 and 28 (purple outline and purple star) cluster together and are separately from the other Fola4 (orange) and Fola1 (blue) clades. The phylogeny was constructed using Geneious Tree Builder using pairwise global alignment with free end gaps and neighbour-joining with *F. solani* as outgroup.

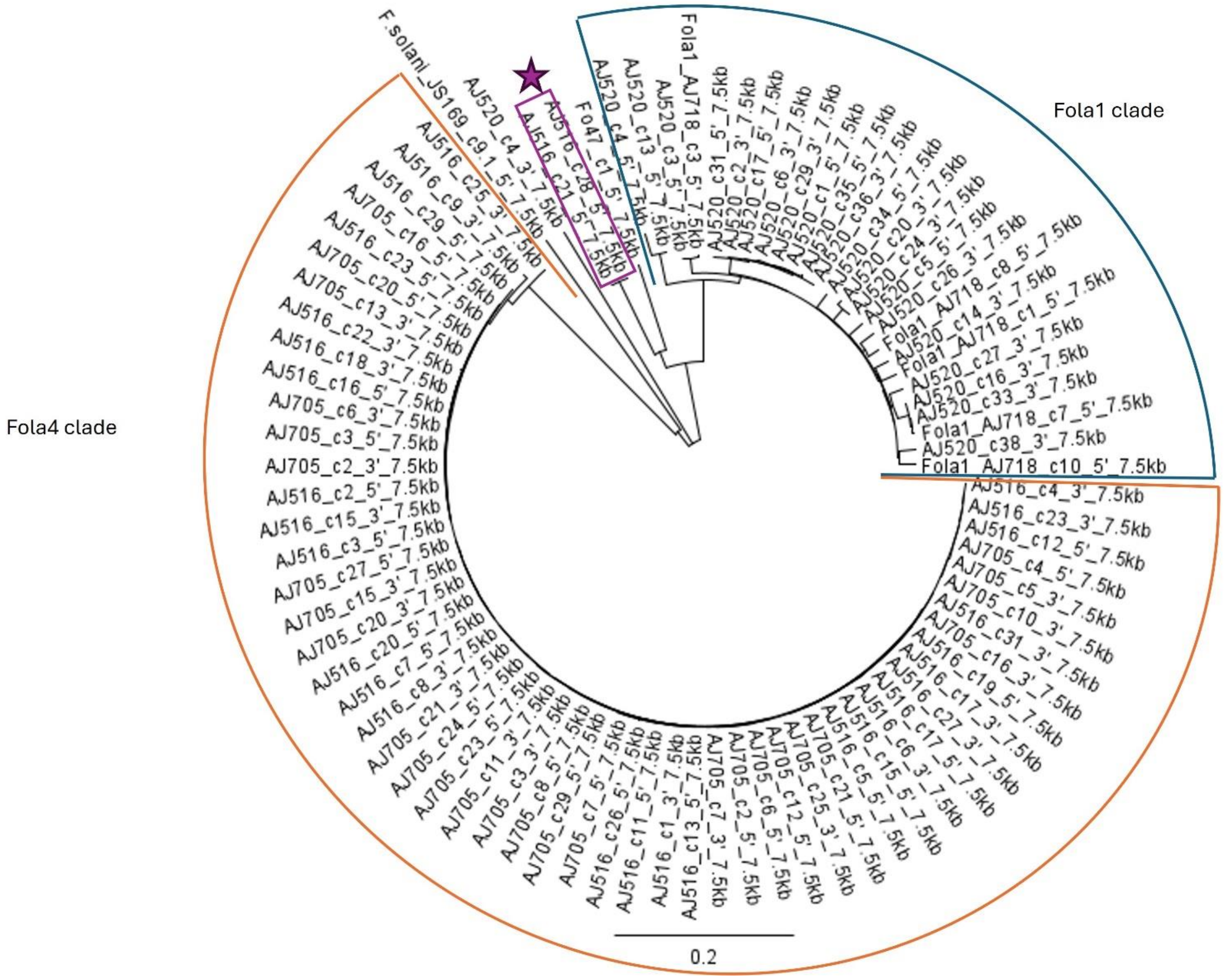
